# Acetylation of conserved lysines fine-tune mitochondrial malate dehydrogenase activity in land plants

**DOI:** 10.1101/2020.10.30.362046

**Authors:** Manuel Balparda, Marlene Elsässer, Mariana B. Badia, Jonas Giese, Anastassia Bovdilova, Meike Hüdig, Lisa Reinmuth, Markus Schwarzländer, Iris Finkemeier, Mareike Schallenberg-Rüdinger, Veronica G. Maurino

**Author notes:** Corresponding authors: Veronica G. Maurino, Mareike Schallenberg-Rüdinger and Iris Finkemeier. These authors contributed equally to the work (Manuel Balparda, Marlene Elsässer, Mariana B. Badia). **Author for contact:** Veronica G. Maurino.

## Abstract

Plants need to rapidly and flexibly adjust their metabolism to changes of their immediate environment. Since this necessity results from the sessile lifestyle of land plants, key mechanisms for orchestrating central metabolic acclimation are likely to have evolved early. Here, we explore the role of lysine acetylation as a posttranslational modification to directly modulate metabolic function. We generated a lysine acetylome of the moss *Physcomitrium patens* and identified 638 lysine acetylation sites, mostly found in mitochondrial and plastidial proteins. A comparison with available angiosperm data pinpointed lysine acetylation as a conserved regulatory strategy in land plants. Focusing on mitochondrial central metabolism, we functionally analyzed acetylation of malate dehydrogenase (mMDH), which acts as a hub of plant metabolic flexibility. In *P. patens* mMDH1, we detected a single acetylated lysine located next to one of the four acetylation sites detected in *Arabidopsis thaliana* mMDH1. We assessed the kinetic behavior of recombinant *A. thaliana* and *P. patens* mMDH1 with site-specifically incorporated acetyl-lysines. Acetylation of *A. thaliana* mMDH1 at K169, K170, and K334 decreases its oxaloacetate reduction activity, while acetylation of *P. patens* mMDH1 at K172 increases this activity. We found modulation of the malate oxidation activity only in *A. thaliana* mMDH1, where acetylation of K334 highly activated it. Comparative homology modelling of MDH proteins revealed that evolutionarily conserved lysines serve as hotspots of acetylation. Our combined analyses indicate lysine acetylation as a common strategy to fine-tune the activity of central metabolic enzymes with likely impact on plant acclimation capacity.

**Significance statement:** We explore the role of lysine acetylation as a mechanism to directly modulate mitochondrial metabolism in land plants by generating the lysine acetylome of the moss *Physcomitrium patens* and comparing with available angiosperm data. We found acetylation of evolutionarily conserved lysines as a strategy to fine-tune the activity of mitochondrial malate dehydrogenase in a species-dependent molecular context.

## Introduction

Flux through central metabolism is particularly dynamic in plants and can change dramatically in response to changes in external conditions or intrinsic demands. For instance, the tricarboxylic acid (TCA) cycle, which is localized in the mitochondria, can operate in a cyclic or non-cyclic mode depending on cell type and the demands for reducing power, ATP, and carbon skeletons (Sweetlove *et al.*, 2010). Although glycolysis-derived pyruvate decarboxylation is considered to be the major entry point for carbon into the TCA cycle in most organisms, in green plant tissues it is malate rather than pyruvate that acts as the main substrate for the TCA cycle under most circumstances (Sweetlove *et al.*, 2010). Mitochondrial malate dehydrogenase (mMDH; EC 1.1.1.37) interconverts malate and oxaloacetate (OAA) using NAD^+^/NADH as co-substrate. *In vivo*, the direction of the reaction depends on the demands of the cells and the redox state of the NAD pool in the matrix (Fig. 1). Apart from its classical role in supporting the TCA cycle flux to generate OAA from malate for respiration, mMDH provides OAA for the synthesis of aspartate, and – indirectly – of citrate as precursor for nitrogen assimilation (Fig. 1) (Hanning and Heldt, 1993, Sweetlove *et al.*, 2010). In particular in illuminated leaves, mMDH can also support metabolic flux in the opposite direction to oxidize NADH to NAD^+^ for consumption by the photorespiratory glycine decarboxylase (Journet *et al.*, 1981), and as part of the mitochondrial malate valve that connects the redox states of the NAD pools of the mitochondrial matrix and the cytosol through the exchange of malate and OAA (Fig. 1) (Scheibe, 2004, Scheibe *et al.*, 2005, Sweetlove *et al.*, 2010). All these aspects of mitochondrial metabolism are supported by models obtained through diel flux balance analysis (Shameer *et al.*, 2019).

**Figure 1.**
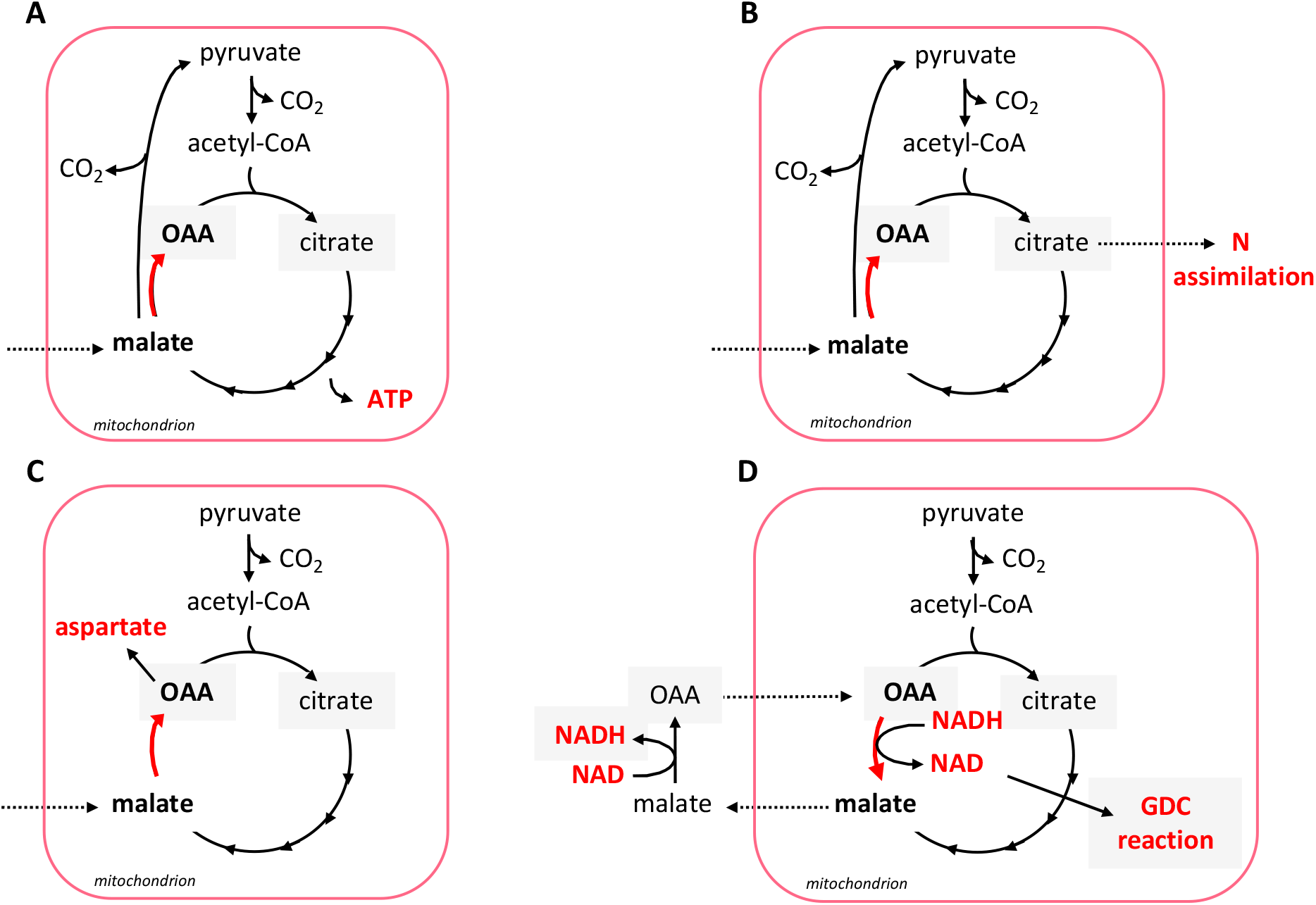
Direction of the mMDH reaction (red arrow) depending on the metabolic demands of the cells and the NAD redox state of the mitochondrial matrix. **A.** Catabolic mode of the TCA cycle. **B.** Provision of OAA for the synthesis of citrate as precursor for nitrogen assimilation. **C.** Provision of OAA for the synthesis of aspartate. **D.** Provision of NAD^+^ for the photorespiratory glycine decarboxylase (GDC) and of malate for redox balance between matrix and cytosol. (Modified from Sweetlove et al., 2010).

Two isoforms, mMDH1 (AT1G53240) and mMDH2 (AT3G15020), are expressed in *A. thaliana,* of which mMDH1 is the main isoform in green tissue (Tomaz *et al.*, 2010, Hüdig *et al.*, 2015). While the *in vivo* mMDH oligomerization state in plants is not yet definitively established, pig heart mMDH is active as a homodimer of ~70 kDa, possessing two equivalent binding sites (Murphey *et al.*, 1967, Noyes *et al.*, 1974, Shore and Chakrabarti, 1976, Gleason *et al.*, 1994). Mitochondrial malate is not only metabolized through mMDH, but also via the NAD-dependent malic enzyme (NAD-ME) (Tronconi *et al.*, 2008, Sweetlove *et al.*, 2010, Maurino and Engqvist, 2015, Tronconi *et al.*, 2020). Both enzymes together provide a remarkable degree of flexibility to plant respiratory metabolism, since they are able to supply mitochondrial carbon metabolism with substrate to respire, but also replenish the TCA cycle with carbon skeletons to maintain its function even when carbon skeletons are withdrawn for biosynthesis, e.g., of amino acids (Sweetlove *et al.*, 2010). While the positioning of mMDH at a branchpoint of central carbon metabolism of plants and its variable engagement depending on flux mode makes active regulation of mMDH activity particularly likely, relatively little is known about this regulation. Modulation of mMDH activity is not only critical to direct carbon flux either to OAA or pyruvate, but it also has the potential to set the total (photo-) respiratory flux by NAD^+^ provisioning to glycine decarboxylase; such modulation can additionally uncouple the redox states of the NAD pools of the matrix and the cytosol (Sweetlove *et al.*, 2010, Shameer *et al.*, 2019). Plant mMDH was not found to be redox-regulated through the matrix thioredoxin system. Instead, it is most probably controlled in response to variations in the matrix adenine nucleotide balance, as its *in vitro* activity is lowered by ATP and inhibited by an increase in the ATP/ADP ratio within the physiological range (Yoshida and Hisabori, 2016). Recently, in *A. thaliana* seedlings growing in liquid cultures and harvested at the beginning of the light period, mMDH1 was found to be acetylated at four different lysine residues: K170, K325, K329, and K334 (König *et al.*, 2014a). However, the functional significance of those modifications has not been explored.

Posttranslational modifications (PTMs), such as phosphorylation and acetylation, can alter protein functions by affecting protein interactions, subcellular localization, or enzymatic activities (Matsuzaki *et al.*, 2005, Yang and Seto, 2008, Ventura *et al.*, 2010, Inuzuka *et al.*, 2012, Bovdilova *et al.*, 2019). Protein acetylation of lysine residues has long been recognized as a regulator of transcriptional control (Allfrey *et al.*, 1964). More recently, this PTM emerged as a regulator of cellular metabolism and signaling in different organisms (Wang *et al.*, 2010, Zhao *et al.*, 2010, Finkemeier *et al.*, 2011, Choudhary *et al.*, 2014, König *et al.*, 2014a).

Lysine acetylation has been found to be particularly widespread in bacterial and mitochondrial proteomes (Xu *et al.*, 2009, Weinert *et al.*, 2011). Two coenzymes of energy metabolism, acetyl-CoA and NAD^+^, are required as substrates for the reversible acetylation of lysine residues (Imai *et al.*, 2000, Lin *et al.*, 2012). Acetyl-CoA is the coenzyme of lysine acetyltransferases but can also acetylate proteins non-enzymatically at a pH higher than eight (Guan and Xiong, 2011, Hirschey *et al.*, 2011, König *et al.*, 2014a). NAD^+^ is used by sirtuin-type deacetylases, which also reside within mitochondria of mammals (Imai *et al.*, 2000) and plants (König *et al.*, 2014b). In mitochondria, acetyl-CoA and NAD^+^ play key roles as metabolic regulators. While acetyl-CoA is produced in the mitochondrial matrix by the pyruvate dehydrogenase complex (PDC) and is oxidized in the TCA cycle, NAD^+^ is required as electron acceptor in the TCA cycle. Respiration rates can have a major impact on protein acetylation, since changes in physiological NAD^+^ concentration correlate with the activity of sirtuins (Lombard *et al.*, 2007, Anderson *et al.*, 2017). The mitochondrial acetylome of 10-day-old *Arabidopsis thaliana* seedlings revealed 120 lysine-acetylated proteins, which contained a total of 243 lysine-acetylated sites (König *et al.*, 2014a). Most TCA cycle enzymes of *A. thaliana* were found to be acetylated, which parallels findings in animals and bacteria (Wang *et al.*, 2010, Masri *et al.*, 2013, König *et al.*, 2014a). That PDC and most of the TCA cycle enzymes were found to be lysine-acetylated across different organisms suggests that acetylation might represent an evolutionary conserved regulation system for TCA cycle function (Hosp *et al.*, 2017).

Lysine acetylation is over-represented in both mitochondria and chloroplasts of angiosperm species, suggesting a prominent role of lysine acetylation in the direct modulation of the function of the endosymbiotic organelles (Hartl *et al.*, 2017, Moller *et al.*, 2020). Little is known about the conservation of acetylated lysines in distant land plant species, as investigations of lysine acetylation hitherto covered only flowering plant species (Melo-Braga *et al.*, 2012, König *et al.*, 2014a, Smith-Hammond *et al.*, 2014, Fang *et al.*, 2015, He *et al.*, 2016, Zhen *et al.*, 2016, Jiang *et al.*, 2018).

Here, we used the model moss *Physcomitrium (Physcomitrella*) *patens* as an early branching land plant (Rensing *et al.*, 2020) to devise a comparative proteomic analysis to shed light on the evolutionary conservation of lysine acetylation in mitochondrial proteins and its potential functional diversification in plants. Our analysis identified lysine acetylation of different mitochondrial proteins of *P. patens,* including mMDH. Using mMDH as a model enzyme of central metabolic importance, we address the question of whether lysine acetylation represents an evolutionary conserved strategy to modulate mMDH activity. We synthesized *A. thaliana* and *P. patens* acetylated mMDH1 proteoforms at the identified lysine acetylation positions using the genetic code expansion strategy (Neumann *et al.*, 2008), and assessed the kinetic behavior of the recombinant enzyme variants. Our results indicate acetylation of conserved lysines as a common strategy to modulate mitochondrial carboxylic acid metabolism by fine tuning mMDH activity.

## Results

### Identification of mitochondrial lysine-acetylated proteins in *P. patens*

To identify lysine-acetylated proteins of the moss *P. patens*, we analyzed its proteome via acetyl-lysine enrichment and liquid chromatography mass spectrometry (LC-MS/MS). The proteome was obtained from gametophores, which represent the haploid and dominant growth stage of mosses, analogous to the sporophyte growth stage of vascular plants (Strotbek *et al.*, 2013). Overall, we detected 6428 protein groups, of which 638 were found to be acetylated on at least one and at most nine lysines (Fig. 2, Supporting Table S1).

**Figure 2:**
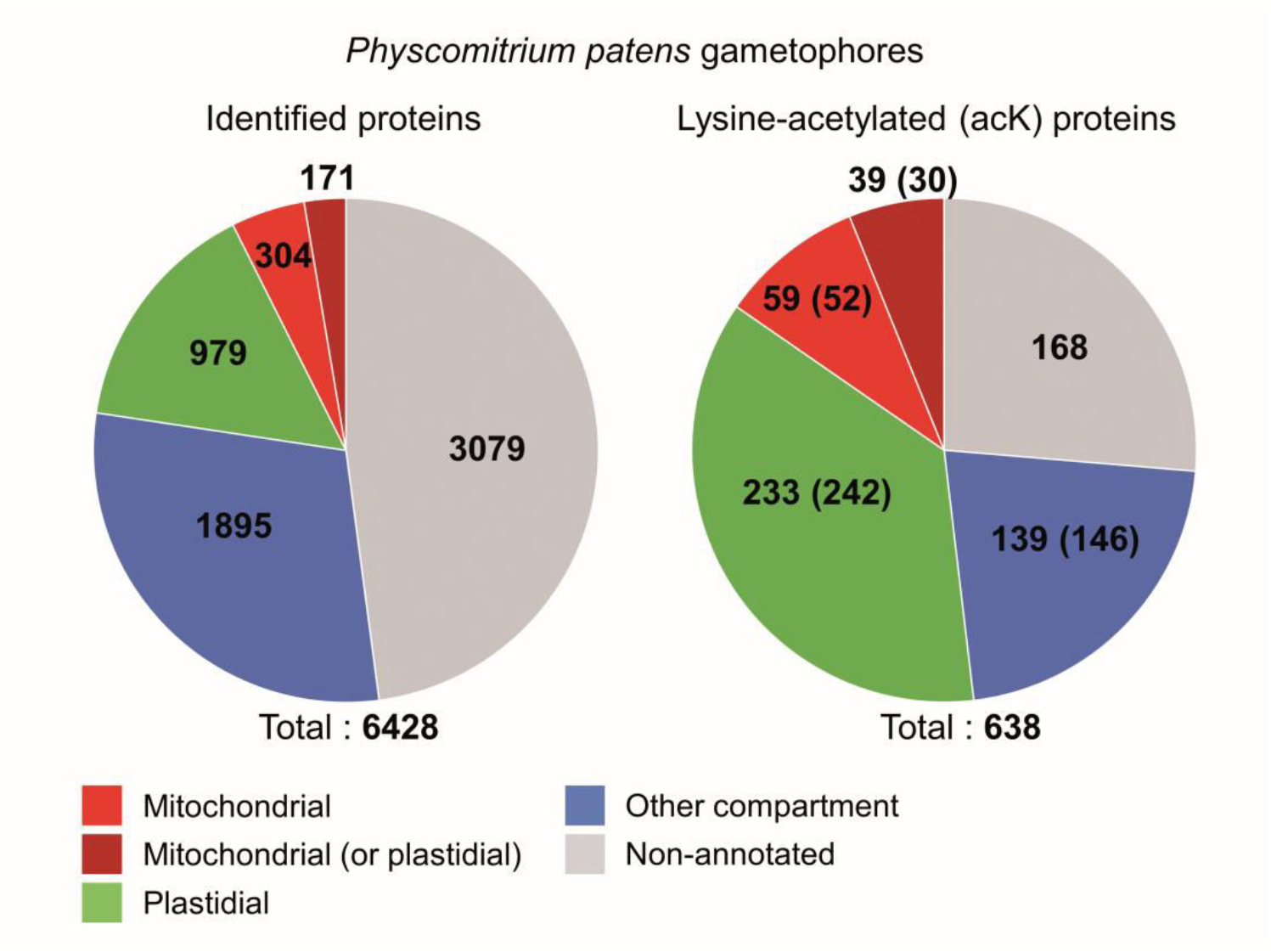
Distribution of lysine acetylation in *P. patens* proteins. Displayed are the proteins identified in gametophore samples of *P. patens* (left) and the proteins with detected acetylation sites (right). The proteins are classified as mitochondrial (red), plastidial (green), other compartment (blue) and non-annotated (grey) based on the localization identified in Müller et al. (2014) and the annotation provided by Lang et al. (2018). Proteins which could only be classified as mitochondrial or plastidial are displayed in dark red. Manually inspection suggested different localisation for 18 proteins; modified numbers are given in brackets. Organellar proteins are overrepresented in the lysine acetylation dataset (p-value <0,0001 for equal distribution of acetylation in organellar proteins and non-organellar proteins, statistical significance was tested with chi-square test using Excel package, XLSTAT (Addinsoft)).

We next classified the identified proteins according to their subcellular localization, making use of a previous analysis of the mitochondrial and plastid proteomes of *P. patens* (Müller *et al.*, 2014). Additional proteins were grouped as organelle-localized when classified as such in the most recent annotation (Lang *et al.*, 2018). In total, 1454 proteins were categorized as organellar proteins, 331 of which were acetylated (Fig. 2). Of those, 272 were annotated as plastidial proteins and 98 as mitochondrial, including 39 proteins that were classified as plastidial and/or mitochondrial (Supporting Table S2 and S3). A post-hoc manual inspection suggested that 16 out of the 98 putative mitochondrial proteins were in fact plastidial (11) or peroxisomal (5; Supporting Table S2), and the putative plastidial citrate synthase proteins were rather peroxisomal due to highest sequence similarity (65-74 %) to the peroxisomal citrate synthases of *A. thaliana*. Correcting these mis-classifications results in 52 mitochondrial, 30 mitochondrial or plastidial, and 242 plastidial proteins identified as lysine acetylated (Fig. 2).

Similar to previous findings in angiosperms (Hartl et al., 2017; Moller et al., 2020), lysine acetylation is particular abundant in the mitochondria and plastids of *P. patens* gametophores. We found that 20% of the identified mitochondrial proteins and 23% of the identified plastidial proteins were acetylated, while only 6 % of the other identified proteins possessed this PTM (Fig. 2; Supporting Table S2). This observation mirrors previous results from isolated mitochondria of *A. thaliana* seedlings, where 120 acetylated protein groups were identified (König *et al.*, 2014a). In *P. patens*, we observed lysine acetylation in most TCA cycle enzymes detected in the dataset (aconitase, NAD- and NADP-dependent isocitrate dehydrogenase, 2-oxoglutarate dehydrogenase, and mMDH), which also mirrors previous findings in *A. thaliana*.

### Different lysine residues are acetylated in TCA cycle enzymes of *A. thaliana* and *P. patens*

The specific sites of lysine acetylation in TCA cycle enzymes and other mitochondrial proteins were compared between orthologous proteins of *P. patens* and *A. thaliana* (König *et al.*, 2014a). Proteins were clustered into orthologous groups, each consisting of one up to six different proteins per species, depending on the number of paralogous proteins encoded in the two genomes and detected in the datasets (Supporting Table S1, Table 1). The PDC E1 α as well as four of the identified TCA cycle enzymes are acetylated in both *P. patens* and *A. thaliana* (Table 1). In contrast, lysine acetylation of NADP isocitrate dehydrogenase proteins was detected in *P. patens* only (Supporting Table S1).

**Table 1.** Conservation of lysine acetylation between mitochondrial TCA cycle and PDC proteins identified in *A. thaliana* and *P. patens.*

| Orthologous protein group | Accession No. | Number of acetylated lysines |  |  |  | Position(s) of acK(s) |
| --- | --- | --- | --- | --- | --- | --- |
|  |  | total | diff | shared K | shared acK |  |
| Pyruvate dehydrogenase complex E1 $\alpha$ | AT1G24180.1 | 2 | 2 | 1 | 1 | <u>85</u>;336 |
|  | AT1G59900.1 | 2 |  |  |  | <u>81</u>;332 |
|  | Pp3c12_7580V3.1 | 1 |  |  |  | <u>86</u> |
|  | Pp3c4_20210V3.1 | 1 |  |  |  | <u>86</u> |
| Aconitase (TCA) | AT2G05710.1 | 3 | 7 | 5 | 1 | <u>790</u>;846;895 |
|  | AT4G26970.1 | 2 |  |  |  | <u>795</u>;900 |
|  | Pp3c15_25170V3.1 | 5 |  |  |  | <u>520</u>;529;598;<u>803</u>;844 |
|  | Pp3c9_25990V3.2 | 2 |  |  |  | <u>491</u>;806 |
| 2-oxoglutarate dehydrogenase E1 (TCA) | AT3G55410.1 | 3 | 5 | 2 | 0 | 549;<u>599</u>;613 |
|  | AT5G65750.1 | 2 |  |  |  | <u>603</u>;873 |
|  | Pp3c23_7750V3.1 | 1 |  |  |  | 1000 |
| NAD isocitrate dehydrogenase (TCA) | AT3G09810.1 | 1 | 3 | 3 | 0 | <u>181</u> |
|  | AT5G03290.1 | 2 |  |  |  | <u>181</u>;218 |
|  | Pp3c15_3920V3.1 | 1 |  |  |  | <u>365</u> |
|  | Pp3c9_4370V3.1 | 1 |  |  |  | <u>355</u> |
| **Malate dehydrogenase (TCA)** | **AT1G53240.1/mMDH H1** | 4 | 5 | 3 | 0 | <u>170</u>;325;329;<u>334</u> |
|  | AT3G15020.1/mMDH2 | 1 |  |  |  | <u>170</u> |
|  | **Pp3c4_20940V3.1/M DH1** | 1 |  |  |  | <u>172</u> |
|  | Pp3c12_8120V3.1/mMDH2 | 1 |  |  |  | <u>170</u> |
| **Sum** | **AT: 10/Pp: 9** | **36** | **22** | **14** | **2** |  |

**Note:** Proteins identified to be lysine-acetylated in *P. patens* and *A. thaliana* are organized into orthologous groups. Total numbers of identified acetylated lysines (K, total), number of acetylated Ks at different positions in the orthologous proteins (diff), numbers of acetylated Ks with conserved lysines at same position of at least one putative ortholog of the other species (shared K) and shared acetylated lysines (shared acK) are listed separately. Positions of the acetylated Ks (acK) in the proteins are given and numbers are underlined, when Ks are conserved in the putative orthologs at this position. Bold numbers represent acetylated lysines, which were detected in at least one *P. patens* and one *A. thaliana* putative ortholog of both datasets. The malate dehydrogenase proteins shown in red are further assessed in this study. Pp: *P. patens*, AT: *A. thaliana*.

Altogether, we found acetylation of 22 different lysines in the TCA cycle enzymes and the E1 α subunit of the PDC in *P. patens* or *A. thaliana*. In 14 of the 22 positions, the lysine, acetylated in one species, was conserved in at least one orthologous protein of the other species. Only the E1 α subunit of the PDC and the aconitase of both species shared a lysine that was acetylated in both datasets (Table 1). A similar picture emerged for the other mitochondrial orthologous protein groups investigated (Supporting Table S4).

Orthologous proteins of *A. thaliana* and *P. patens* showed conserved lysine positions, but for each individual position, lysine acetylation was typically detected in one of the two species only (Table 1, Supporting Table S4).

In addition to the TCA cycle enzymes and the PDC, we classified 25 mitochondrial *A. thaliana* proteins and 31 mitochondrial *P. patens* proteins into 17 further orthologous groups. We found 99 lysine residues to be acetylated in at least one protein of these additional orthologous groups. 66 of these 99 positions (67%) carried a lysine in orthologous proteins of the respective other species as well; 12 (12%) of these conserved lysines were acetylated in orthologous proteins of both species (Supporting Table S4). Mitochondrial orthologous proteins of *A. thaliana* and *P. patens* showed 66% conserved lysine positions (80 positions/121 positions), which were found to be acetylated in at least one of the orthologs compared in this study. For each individual position, however, lysine acetylation was typically detected in one of the two species only (Table 1, Supporting Table S4).

*A. thaliana* mMDH was found acetylated on four lysines (K170, K325, K329, and K334) (König *et al.*, 2014a), while *P. patens* mMDH was acetylated in one unique lysine (K172 of Pp3c4_20940V3.1, in the following named mMDH1, or K170 of Pp3c12_8120V3.1, in the following named mMDH2) corresponding to K169 of *A. thaliana* mMDH1 (Table 1, Fig. 3). Acetylation at K170 of *A. thaliana* mMDH proteins and K172 of *P. patens* mMDH proteins could not be unequivocally assigned to one of the two paralogs, mMDH1 and mMDH2, present in each species (Supporting Fig. S1). However, in the proteomic datasets of both *A. thaliana* and *P. patens*, mMDH1 was more abundant than mMDH2 (Supporting Table S1, König et al. 2014). Hence, we focused our further analyses on the dominant mMDH protein, mMDH1, of both species.

**Figure 3.**
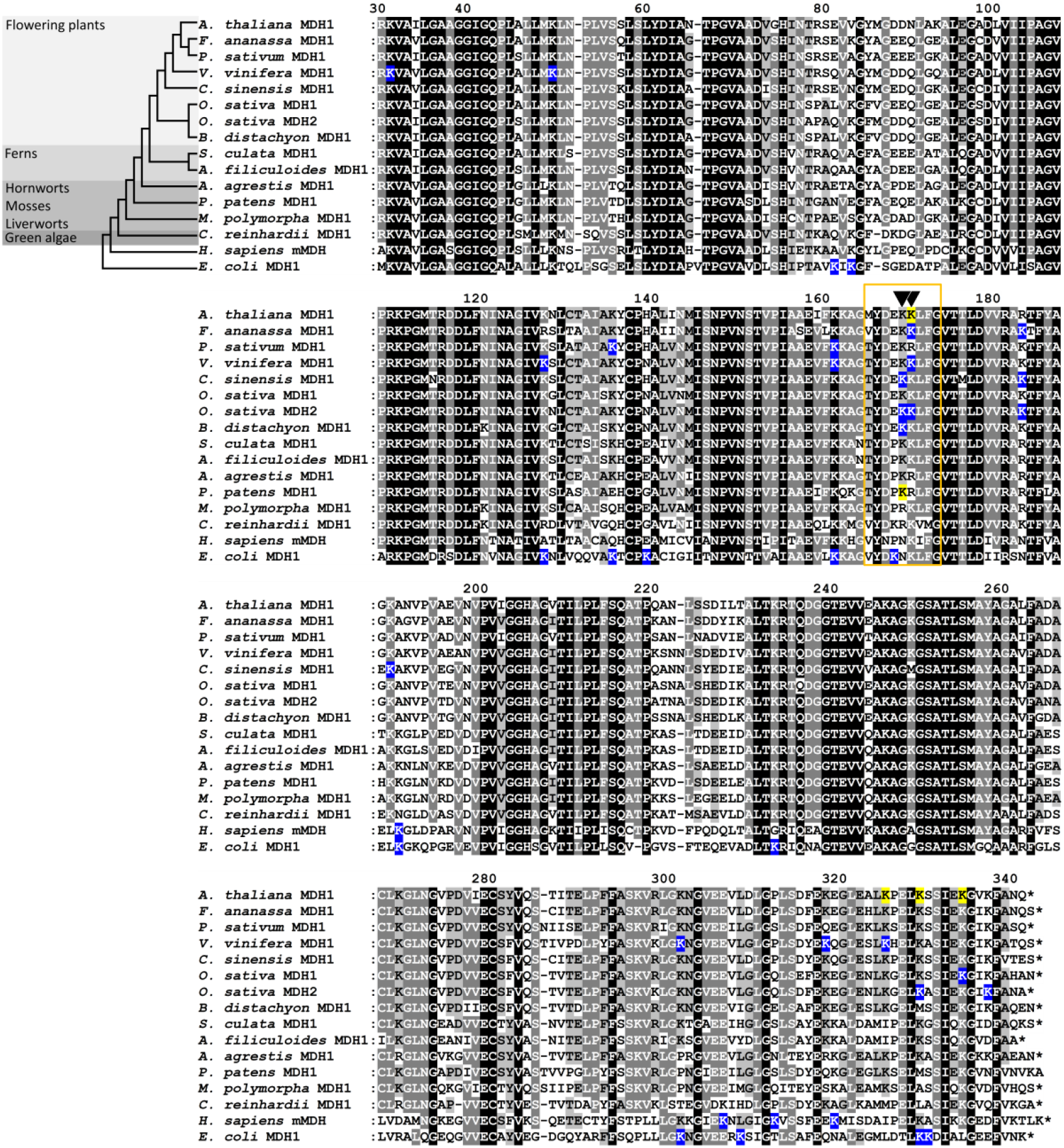
Sequence conservation of mMDH of representative species of each main land plant clade, human and *E. coli*, and their lysine acetylation sites. Alignment of the acetylated protein isoforms of the mMDHs of the angiosperms *A. thaliana*, *F. ananassa*, *P. sativum*, *V. vinifera*, *C. sinensis*, *O. sativa* and *B. distachyon* with found lysine acetylation sites marked in blue. Orthologues of representative species of other major land plant clades (the ferns *A. filiculoides* and *S. cucullata,* the hornwort *A. agrestis,* the moss *P. patens*, the liverwort *M. polymorpha* and the green algae *C. reinhardtii*) share high sequence conservation. Shading in black indicates 100%, in dark grey >80% and in light grey >60% amino acid identity conservation. The cladogram is based on the current understanding of land plant phylogeny (Qiu *et al.*, 2006, Bremer *et al.*, 2009). The sequences of *H. sapiens* mitochondrial MDH2 and *E. coli* MDH with their identified acetylated lysines (blue) are included for comparison. The numbering is based on the sequence of MDH1 of *A. thaliana*. The alignment starts from amino acid 30 which corresponds to the start methionine of *E. coli* MDH. Yellow highlighted Ks are the acetylated lysines in *A. thaliana* and *P. patens* mMDH1 analyzed in this study. A region with acetylation sites conserved in different species is framed in orange and includes position K169 and K170 marked with black arrows.

### Lysine acetylation of mMDH along the phylogeny of land plants

To analyse if acetylation of mMDH is evolutionarily conserved across land plants, we inspected available acetylome datasets of nine additional flowering plant species for the presence of acetylation sites in mMDH (Supporting Table S5). In six of these species, unique acetylated peptides of mMDHs were detected (Fig. 3, Supporting Fig. S1, Supporting Table S5). The species with available information on mMDH acetylation cover six different orders of flowering plants (*A. thaliana*: Brassicales (König et al. 2014), *Fragaria ananassa*: Rosales (Fang *et al.*, 2015), *Pisum sativum:* Fabales (Smith-Hammond et al. 2014), *Vitis vinifera:* Vitales (Melo-Braga *et al.*, 2012), *Camellia sinensis:* Ericales (Jiang et al. 2018), *Oryza sativa* and *Brachypodium distachyon:* Poales (Zhang *et al.*, 2015, He *et al.*, 2016, Xiong *et al.*, 2016, Zhen *et al.*, 2016, Zhou *et al.*, 2018)).

We aligned the mMDH protein sequences of these six species with mMDHs of model plants representing different main land plant clades (Ferns: *Azolla filiculoides* and *Salvinia cucullata*; hornworts: *Anthoceros agrestis*, liverworts: *Marchantia polymorpha* and green algae: *Chlamydomonas reinhardtii*) including *P. patens* as model moss and *A. thaliana* as model flowering plant. Sequences were identified via Blast using *A. thaliana* mMDH1 as query. For selected model species, we identified MDH proteins clustering together with mMDH of *A. thaliana* and *P. patens* in phylogenetic analyses (Supporting Fig. S1). Peroxisomal and chloroplastic MDHs of *A thaliana* and *P. patens* clustered in separate clades. For comparison, we included the *E. coli* MDH and human mMDH2 along with their known lysine acetylation sites (Zhao *et al.*, 2010, Schilling *et al.*, 2015) in the sequence alignment (Fig. 3). Sequences in the alignment are clustered based on the land plant phylogeny (Qiu *et al.*, 2006, Bremer *et al.*, 2009). Only for *Oryza sativa*, different acetylation sites in both mMDH paralogs were identified, and thus both proteins were included in the alignment (Zhao *et al.*, 2010, He *et al.*, 2016) (Fig. 3).

An interesting pattern emerges for the two neighboring lysine residues in the mMDH sequences that correspond to K169 and K170 in the *A. thaliana* sequence (Fig. 3). In different studies, these lysine residues were found acetylated in mMDHs of several plant species. K172 of *P. patens* mMDH1 (corresponding to K169 of *A. thaliana*), identified with high coverage in our *P. patens* acetylome, was also acetylated in mMDH1 of *Camellia* (Jiang *et al.*, 2018) and *Brachypodium* (Zhen *et al.*, 2016) leaves, and in mMDH2 of *O. sativa* leaves (Zhou *et al.*, 2018). K170 of *A. thaliana* mMDH1 was also acetylated in mMDH1 of strawberry leaves (Fang *et al.*, 2015) and grape vine exocarp (Melo-Braga *et al.*, 2012), and in mMDH2 of *O. sativa* embryos (He *et al.*, 2016) (Fig. 3). In rice mMDH2, the lysines at position 169 and 170 were identified as differentially acetylated in different organs. The other angiosperms in this dataset showed acetylation only on one or the other lysine (Fig. 3).

Interestingly, at positions corresponding to K169 and K170 in *A. thaliana*, all other investigated species show either a lysine or an arginine; the latter is similar in structure and chemical properties to the non-acetylated lysine. Even in the plastidial and peroxisomal MDHs of *A. thaliana* and *P. patens*, either a lysine or an arginine is present at these two positions (Supporting Table S4). The peroxisomal MDH1 of *A. thaliana* shares the acetylation site with mMDH1 of *P. patens* (Supporting Fig. S1, Supporting Table S4, (König *et al.*, 2014a)). In the *E. coli* MDH and human MDH, however, position 169 is occupied by an asparagine, which would partially mimic the constantly acetylated lysine. Other acetylated lysines in *A. thaliana* mMDH1 are K325, K329, and K334 (König *et al.*, 2014a), each of which was found to be acetylated in at least one other angiosperm species. Acetylated K325 was detected in *V. vinifera*, while acetylated K329 and K334 were found in *O. sativa* mMDH2 and mMDH1, respectively (Fig. 3). In addition, K329 was identified to be acetylated in *E. coli* MDH (Fig. 3).

### Enzymatic properties of site-specific acetylated *A. thaliana* and *P. patens* mMDH1 proteoforms

The conservation of lysine acetylation sites in plant mMDH (Fig. 3) prompted us to investigate the influence of acetylation on the enzymatic properties of the enzyme in the model plants *A. thaliana* and *P. patens*. We selectively incorporate acetyl-lysine (AcK) at the lysines K170, K325, K329, and K334 of mMDH1 of *A. thaliana* (König *et al.*, 2014a) as well as K172 of mMDH1 of *P. patens* (Table 1, Fig. 3, Supporting Tables S2 and S3). The lysine acetylation was incorporated into recombinantly expressed proteins using the amber suppression system in *E. coli*, which allows the co-translational addition of N-acetyl lysine in response to a stop codon at the desired positions (Neumann *et al.*, 2008, Neumann *et al.*, 2009). *A. thaliana* and *P. patens* non-modified mMDH1 and the protein versions carrying the single acetylated lysines were expressed in *E. coli* in the presence of N-acetyl lysine and nicotinamide (to inhibit the *E. coli* deacetylase CobB) and purified to homogeneity (Fig. 4A). In all cases, we obtained proteins of the expected molecular masses (33.5 kDa for *A. thaliana* and 36 kDa for *P. patens*; Fig. 4A). The incorporation of AcK at the desired positions of *A. thaliana* and *P. patens* mMDH1 was confirmed by LC-MS/MS (Supporting Table S6).

**Figure 4.**
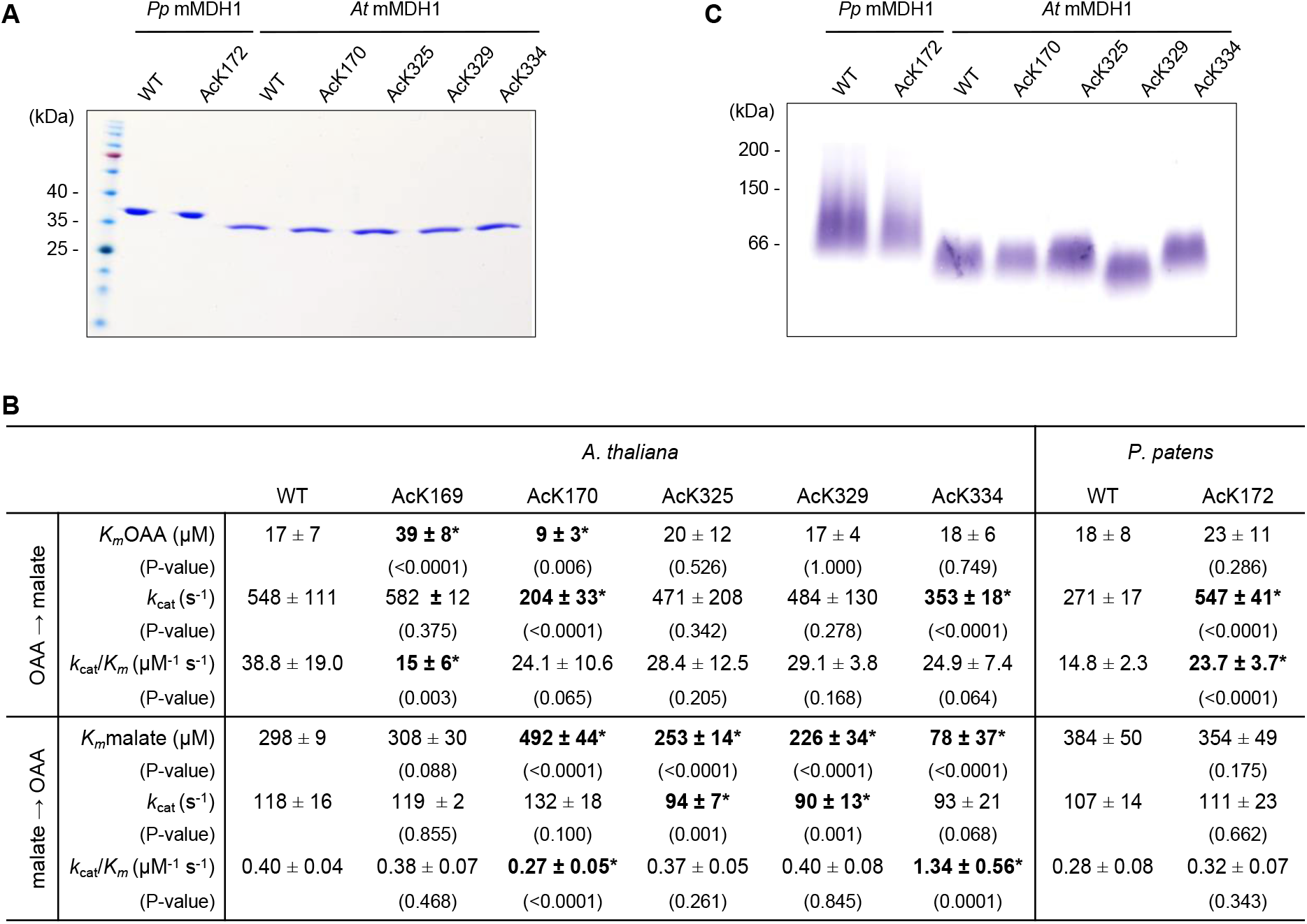
The impact of lysine acetylation on mMDH activity. **A.** SDS-PAGE stained with Coomassie of the isolated recombinant non-modified (denoted as WT) and lysine acetylated (AcK) variants of *P. patens* (*Pp*) and *A. thaliana* (*At*) mMDH1. To the left, molecular weight markers (Spectra Multicolor Broad Range Protein Ladder; ThermoFisher Scientific). **B.** Kinetics parameters of the reduction of OAA and oxidation of malate by recombinant non-modified (denoted as WT) and AcK versions of *A. thaliana* and *P. patens* mMDH1. Data were adjusted to Michaelis-Menten equation by non-lineal regression with Prism 6 (GraphPad Software). The values represent the mean ± standard deviation; n = at least three independent enzyme preparations, each measured in triplicate. **\*** denotes values that are are statistically significantly different from the corresponding WT evaluated by the unpaired t-test. The p-values obtained are indicated in brackets under the evaluated values. **C.** Native PAGE of recombinant non-modified (WT) and lysine-acetylated (AcK) variants of *P. patens* (*Pp*) and *A. thaliana* (*At*) mMDH1. The position of the molecular weight markers (Sigma Aldrich) ß-amlyase (200 kDa), alcohol dehydrogenase (150 kDa) and bovine serum albumin (66 kDa) are shown in the left.

We next determined the *in vitro* kinetic parameters of the purified recombinant non-modified mMDH1 and the single-site acetylated versions. First, we focused on the direction of the reduction of OAA, in which case mMDH activity is needed predominantly to produce malate for its translocation to the cytosol for redox balance (Fig. 1D). We found that acetylation at positions K325 and K329 in *A. thaliana* mMDH1 had no significant effects on the kinetic parameters of the enzyme. In contrast, acetylation at K334 decreased the turnover number (*k*_cat_) to 64.4 % of the non-modified enzyme. Acetylation at K170 decreased the turnover number to 37% while increasing the affinity for OAA (decreasing *K_m_*) to 50 % of the non-modified enzyme. The same analysis with *P. patens* mMDH1 indicated that the acetylation in K172 doubles the turnover number in comparison to the non-modified enzyme (Fig. 4B), with a corresponding increase in the catalytic efficiency (*k*_cat_/*K_m_* = 23.7 ± 3.7 μM^−1^ s^−1^) compared to the non-modified enzyme (*K*_cat_/*K_m_* = 14.8 ± 2.3 μM^−1^ s^−1^).

We further analysed mMDH activity in its respiratory role in the TCA cycle, in which OAA is generated by oxidation of malate (Fig. 1). We found that the acetylation of all analyzed single amino acid positions in *A. thaliana* mMDH1 affected the enzymatic parameters. Acetylation at position K170 reduces the affinity for malate to 65 % compared to the non-modified enzyme (Fig. 4B). As a consequence, K170 acetylation reduces the catalytic efficiency (*K*_cat_/*K_m_* = 0.27 ± 0.05 μM^−1^ s^−1^) to 67.5 % of the non-modified enzyme (*K*_cat_/*K_m_* = 0.40 ± 0.04 μM^−1^ s^−1^). Acetylation at positions K325, K329, and K334 in *A. thaliana* mMDH1 reduced the turnover number to 76-79 % compared to the non-modified enzyme, while increasing the affinity for malate (reduced *K_m_* values, Fig. 4B), by 15 %, 24 %, and 74 %, respectively, compared to the non-modified enzyme (Fig. 4B). The changes in the kinetic parameters of acetylated mMDH1 at K334 result in a 3-fold enhancement of the catalytic efficiency (*K*_cat_/*K_m_* = 1.34 ± 0.56 μM^−1^ s^−1^). In contrast, the catalytic efficiencies of the other two enzymatic variants bearing acetylation at K325 (*K*_cat_/*K_m_* = 0.37 ± 0.05 μM^−1^ s^−1^) and at K329 (*K*_cat_/*K_m_* = 0.40 ± 0.08 μM^−1^ s^−1^) were similar to that of the non-modified enzyme. Acetylation of *P. patens* mMDH1 in K172 had no effects on the enzymatic parameters in the malate oxidation direction compared to the non-modified enzyme (Fig. 4B).

Lysine residues corresponding to K172 of MDH1 of *P. patens* were also found acetylated in three angiosperm species (Zhen *et al.*, 2016, Jiang *et al.*, 2018, Zhou *et al.*, 2018). We further investigated the potential conserved regulatory role of this lysine residue in *A. thaliana* mMDH1. We produced the recombinant *A. thaliana* mMDH mutant AcK169 (corresponding to K172 in *P. patens*) and evaluated its kinetics parameters in both directions of the reaction. We found that, similar to *P. patens,* the acetylation of Arabidopsis mMDH1 in K169 had no significant effects on the kinetic parameters of the enzyme in the malate oxidation reaction. In contrast, in the direction of the reduction of OAA, acetylation at K169 decreases the affinity of Arabidopsis mMDH1 for OAA by 2.3-fold (*K_m_* = 39 ± 8 μM^−1^; p = <0,0001), with a corresponding decrease in the catalytic efficiency (*K*_cat_/*K_m_* = 15 ± 6 μM^−1^ s^−1^; p = 0,0027) compared to the non-modified enzyme (Fig. 4B).

### Mobility analysis of the acetylated proteins by native PAGE

The introduction of a PTM can cause changes in structural features of a protein, which can impact the kinetic properties. To analyse if the general organization of the non-modified enzymes is maintained in the single acetylated mMDH recombinant variants produced, we compared the mobility of the enzymes in native gel electrophoresis. The isolated proteins were run side-by-side on the same gel and analyzed for MDH activity.

We found that in homogeneous native PAGE, the recombinant mMDH1 of *A. thaliana* and *P. patens* are present as single bands very likely corresponding to dimers (Fig. 4D). *A. thaliana* and *P. patens* mMDH1 proteins differ slightly in their mobility, which is likely due to the differences in the molecular masses of the subunits (Fig. 4A) and of different gross shapes of the proteins that can be adopted in solution. The native PAGE results demonstrate that all acetylated mMDH1 versions of *A. thaliana* and *P. patens* conserved the same oligomeric arrangements observed in the non-modified proteins (Fig. 4C), indicating that the detected activity changes cannot be accounted for by changes in homomerisation.

### Mapping of acetylation sites on the crystal structures of MDH

We used the crystal structures of *E. coli* MDH (PDB ID:1EMD) and human MDH2 (PDB ID: 2DFD) to perform homology modelling of the *A. thaliana* and *P. patens* structures and to map the acetylated lysine residues analysed in this study onto the molecular structures. Models of the *A. thaliana* and *P. patens* mMDH1 monomers, dimers, and tetramers show that the acetylated lysine residues are neither located in the direct vicinity of the active site (Fig. 5A-B) nor of the dimer or tetramer interfaces predicted by the constructed models (Fig. 5C-F).

**Figure 5.**
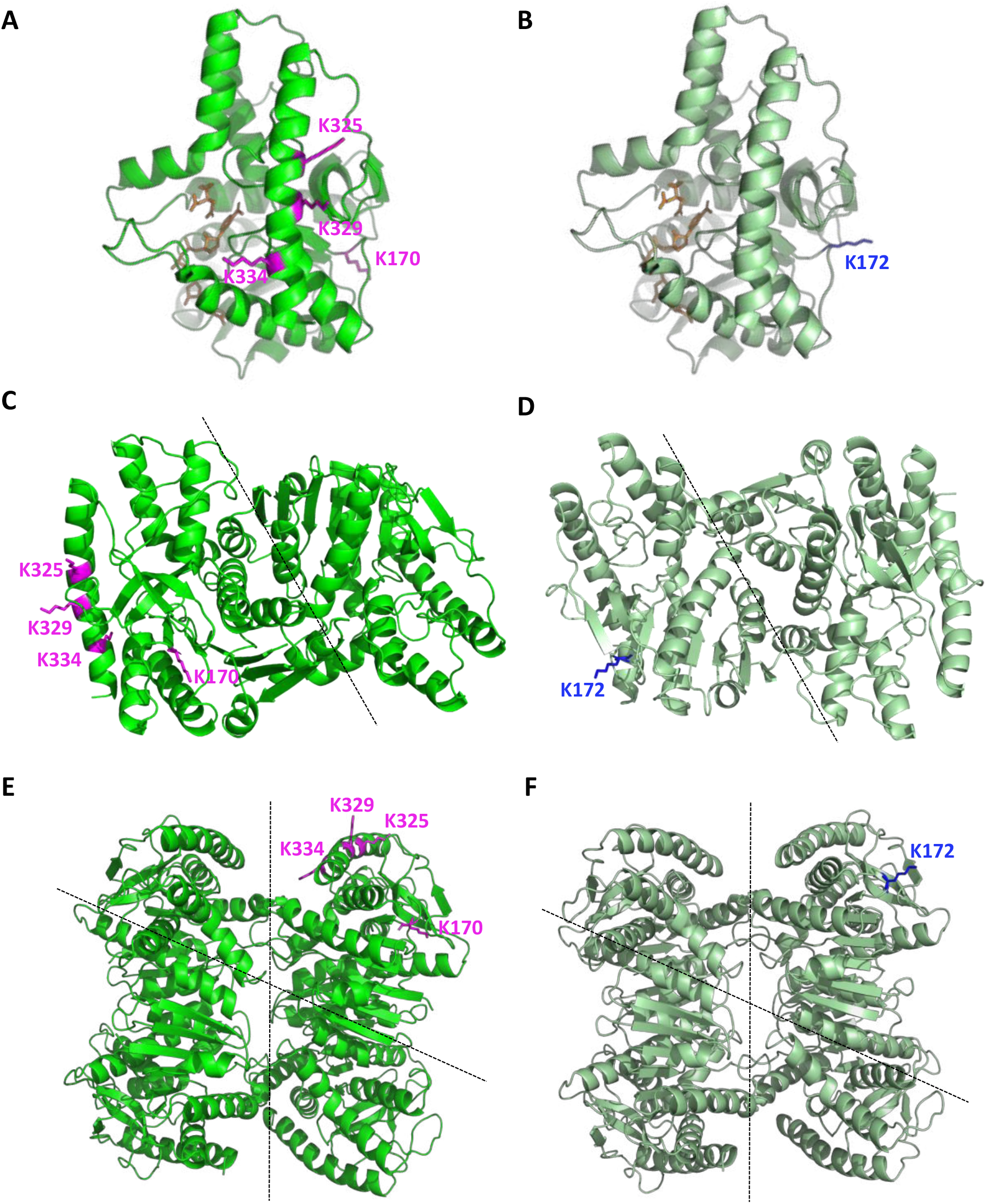
Structures of *A. thaliana* and *P. patens* mMDH1 as obtained by homology modelling. Structures of *A. thaliana* (bright green **A, C, E**) and *P. patens* (dark green; **B, D, F**) mMDH1 modelled as monomers (A and B), dimers (B and D), and tetramers (E and F). The structures were modelled using the crystal structure of *E. coli* MDH (PDB ID:1EMD) and human MDH2 (PDB ID: 2DFD). The acetylated lysine residues of *A. thaliana* mMDH1 are shown in magenta. The acetylated lysine residue of *P. patens* mMDH1 is shown in blue. In A and B, the ligand molecules NAD^+^ and citrate - in brown - were manually fitted in by aligning the structure of the *H. sapiens* MDH2. Dotted lines indicate the dimer and tetramer interfaces.

Superposition analysis of the monomeric structures indicate that *A. thaliana* K170, *P. patens* K172, and *E. coli* K140 are located in the same region of the three-dimensional structure of the proteins (Fig. 6A-C), suggesting conservation of the regulatory function of these lysines by acetylation. The specific spatial orientation of the lysine side chains may differ from the orientations observed in the structures of *A. thaliana* and *P. patens*, however, since the shown structures are modelled based on the crystal structures of *E. coli* and human MDH.

**Figure 6.**
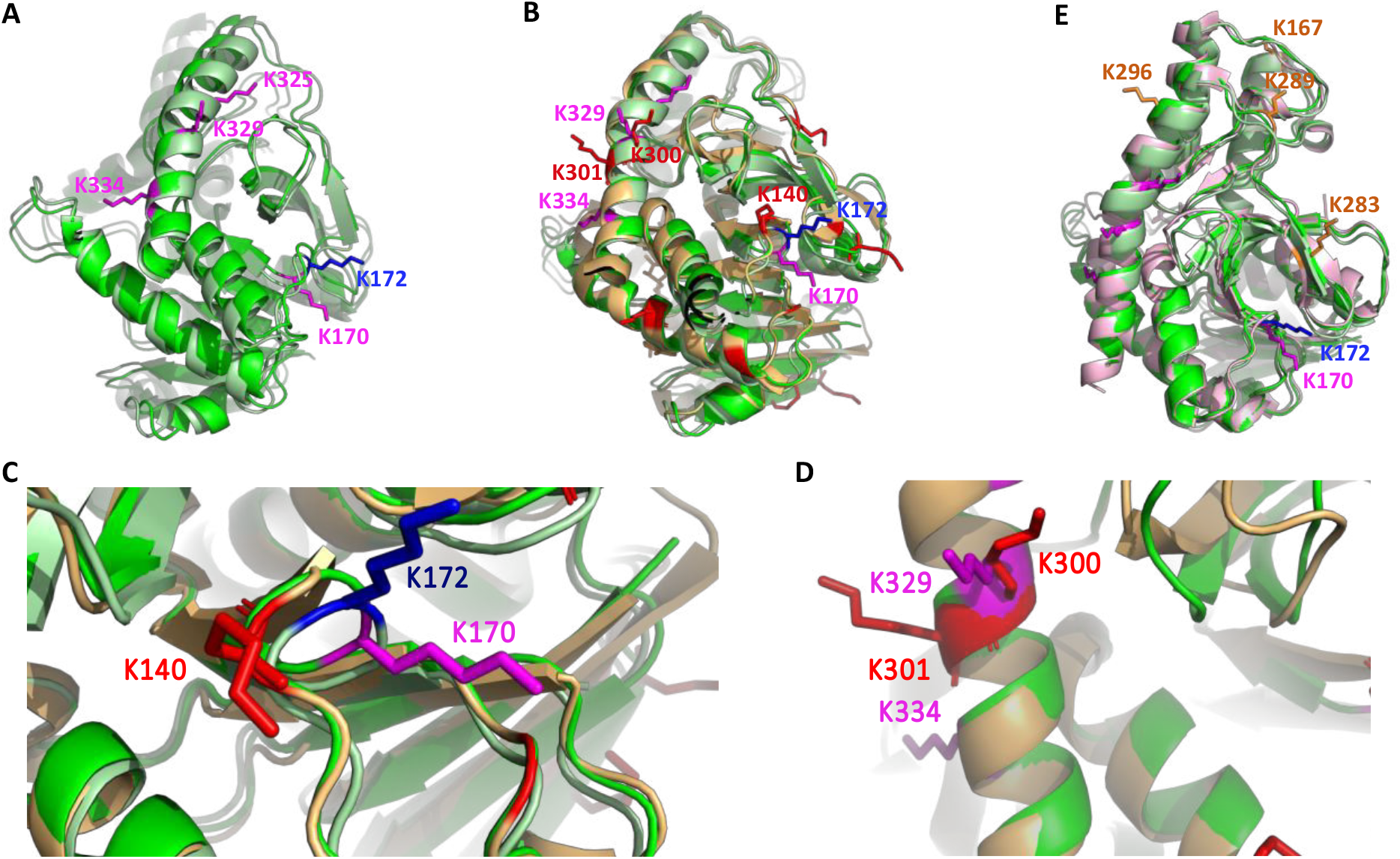
Superposition analysis of MDH monomeric structures. **A**. Structures of *A. thaliana* (bright green-magenta) and *P. patens* (dark green-blue) mMDH1. **B**. Structures of *A. thaliana* and *P. patens* mMDH1, and *E. coli* MDH (gold-red; PDB ID:1EMD). **C.** Zoom in of the region containing K170 in *A. thaliana* (bright green-magenta) mMDH1, K172 in *P. patens* (dark green-blue) mMDH1, and K140 in *E. coli* (gold-red) MDH. **D.** Zoom in of the carboxy-terminal alpha helix containing K329 and K334 in *A. thaliana* (bright green-magenta) mMDH1 and K300 and K301 in *E. coli* (gold-red) MDH. **E.** *A. thaliana* and *P. patens* mMDH1, and human MDH2 (rose-brown; PDB ID: 2DFD). The acetylated lysine residues are shown in magenta for *A. thaliana* mMDH1, blue for *P. patens* mMDH1 red for *E. coli* MDH, and brown for human MDH2. For simplicity not all lysines found to be acetylated in *E. coli* are numbered.

We also observed that *A. thaliana* mMDH1 K329 and K334, two lysine positions that are conserved in plants, are located in the same amino-terminal alpha-helix as K300 and K301 of *E. coli* MDH (Fig. 6B, D), with K329 of *A. thaliana* mMDH1 occupying the same special position as K300 of *E. coli* MDH (Fig. 6B, D). While the effects of acetylation of K300 and K301 on the activity of *E. coli* MDH have not yet been investigated, the conserved acetylation of K300 within plants suggests a possible regulatory function. The superposition of the structures indicates that all acetylated lysine residues of *A. thaliana* and *P. patens* mMDH1 are located in different regions of the protein surface than those of human mMDH2 (Fig. 6E), anticipating different regulation strategies in plants and mammals.

## Discussion

### Lysine acetylation in mitochondrial proteins of evolutionarily distant species

Acetylation of lysine residues is a particularly abundant posttranslational modification of mitochondrial and plastid proteomes (Lombard *et al.*, 2007, König *et al.*, 2014a, Smith-Hammond *et al.*, 2014, Hartl *et al.*, 2017). We detected 324 lysine acetylation sites in organellar proteins of *P. patens* gametophores. Since lysine acetylation is a highly dynamic modification and its occurrence depends on the metabolic status of the organism (Meyer *et al.*, 2018), the acetylation pattern may differ across developmental stages, organs, and growth conditions. To reduce the influence of such intra-organismic variation, we aimed at considering analogous tissues for the comparison between *A. thaliana* and *P. patens*. Mitochondrial proteins were particularly overrepresented amongst lysine acetylated proteins detected in *P. patens*, even more so than in recent whole tissue acetylome studies of *A. thaliana* (Hartl et al. 2017, Uhrig et al. 2019). The number of 82 mitochondrial proteins identified here from whole *P. patens* gametophores is overall in line with 120 lysine-acetylated proteins that were detected in isolated *A. thaliana* mitochondria of seedlings (König *et al.*, 2014a).

Due to the lack of evidence for a mitochondrial acetyltransferase in plants, it appears likely that mitochondrial protein acetylation mostly occurs non-enzymatically between acetyl-CoA and specific lysine residues of particularly high reactivity. The functional steady-state lysine acetylation pattern is likely set by enzymatically controlled de-acetylation, as carried out by sirtuin proteins (Lombard *et al.*, 2007, König *et al.*, 2014b, Anderson *et al.*, 2017). In general, lysine acetylation removes the positive charge from the lysine side chains of proteins, increasing their hydrophobicity. However, similar to phosphorylation, lysine acetylation can affect protein function in various ways, and it can cause activation as well as inhibition of enzymes depending on the function of the particular lysine residue within the protein structure (Hosp *et al.*, 2017).

Although the same mitochondrial proteins are frequently found acetylated in different angiosperm species, the location of the particular acetylated lysines within the protein often differs (Finkemeier *et al.*, 2011, Hosp *et al.*, 2017). Our study shows that this is true even across land plant phylogeny. We detected the same mitochondrial proteins acetylated in the distant land plant species *P. patens* and *A. thaliana* (in total 22 orthologous protein groups were identified) and found that the majority (66%) of lysine positions that were acetylated in mitochondrial proteins of *A. thaliana* or *P. patens* was conserved in the orthologs of the other species as well (Supporting Table S4). However, even in the TCA cycle or PDC proteins, which showed high acetylation in both *P. patens* and *A. thaliana* (up to 4 acetylated lysines), only a low number of acetylation sites were shared (in total two) between the orthologs of the two species. A reason for the limited number of shared acetylated lysine residues detected might be the pronounced dynamics in lysine (de-) acetylation, such that each experimental condition may only detect a subset of physiologically relevant acetylations. The enzymes of the TCA cycle are known to be regulated by reversible acetylation in bacterial and mammalian cells, in dependance on the nutritional status as well as on the circadian clock (Wang *et al.*, 2010, Masri *et al.*, 2013, Meyer *et al.*, 2018). Whereas caloric restrictions generally lead to deacetylation, nutrition with sugars leads to a strong increase in mitochondrial acetylation of all TCA cycle enzymes (Meyer *et al.*, 2018).

Lysine residues corresponding to K170 in *A. thaliana* mMDH1, for which acetylation led to changes in the enzyme catalytic parameters, were found acetylated in three other species and also under different conditions. The same was observed for site K172 of MDH1 of *P. patens*; the corresponding lysine residues were also found acetylated in three angiosperm species. The detection of homologous acetylated lysines in distant species highlights those PTMs most likely to play a physiologically relevant role in metabolic regulation.

### Acetylation of lysines within a conserved hotspot modulates plant MDH activity

In addition to the identification of acetylation of *E. coli* MDH and human mMDH2 (Venkat *et al.*, 2017), recent proteomic studies report acetylation of plant mMDH proteins, albeit in different sequence positions (Supporting Table S5). To understand the potential functional significance of mMDH acetylation, we examined the enzyme characteristics in both directions of the reversible enzymatic reaction.

Our analysis of the kinetic properties of plant mMDH1 in the OAA reduction direction indicated that lysine acetylation of K172 in *P. patens*, and K169 (corresponding to K172 in *P. patens*) and K170 in *A. thaliana* have species-dependent opposite effects on the activities of the enzyme. Acetylation at K172 of *P. patens* mMDH1 enables the enzyme to convert OAA into pyruvate with a doubled catalytic rate compared with the non-modified protein. In contrast, the incorporation of an acetyl group at K169 or K170 in *A. thaliana* mMDH1 gives rise to less efficient enzymes; acetylation of K169 decreases the affinity for OAA by 2-fold without modifying the catalytic rate, while acetylation of K170 decreases the catalytic activity by just above a third of the rate compared with the non-modified protein. A recent analysis of the protein composition of an individual average plant mitochondrion from cultured heterotrophic *A. thaliana* cells indicated that mMDH1 is highly abundant, with about 13.190 copies per mitochondrion (Fuchs *et al.*, 2020). Adding the *in vitro* activity changes measured here to this model, the non-modified mMDH1would have the capacity to turn over 7.2 Mio molecules of OAA per second in a single mitochondrion, while acetylation at K170 would reduce this conversion capacity to 2.7 Mio molecules per second.

We found that the turnover number of *A. thaliana* non-modified mMDH1 is twice as high as that of *P. patens* non-modified mMDH1 (Fig. 4B), suggesting that the enzymes differ in their basal properties toward the substrates in the different species. These differences are likely to appear as adaptations to the metabolic environments in the species and probably result from adaptive configurations at the active site and/or of more distal regions of the proteins. Interestingly, the opposite effects of acetylation of the lysine residues in both enzymes change the turnover numbers of the acetylated proteins to values similar to those found in the non-modified enzyme of the other species analysed: the turnover number of *P. patens* mMDH1 acetylated in K172 has a similar value to that of *A. thaliana* non-modified mMDH1, while *A. thaliana* mMDH1 acetylated in K170 has a similar turnover number to that of *P. patens* non-modified mMDH1. These observations suggest that changes of the turnover number induced by acetylation of plant mMDH1 occur between minimal and maximal values that are set by the structure and catalytic mechanism of the plant enzyme. Although the precise molecular mechanism for the changes in the turnover number are still to be determined, it is interesting to note that amino acid 173 in *P. patens* mMDH1 (corresponding to K170 in *A. thaliana* mMDH1) is an arginine, which has similar properties as a permanent non-acetylated lysine (Kamieniarz and Schneider, 2009). Considering that the presence of an acetyl-lysine at position 170 of *A. thaliana* mMDH1 decreases the turnover number of the enzyme, the presence of an arginine in *P. patens* mMDH1 at the same position suggests that *P. patens* mMDH1 is adapted to avoid a further reduction of its enzymatic activity, at least by acetylation. This interpretation is in accordance with the lower *k*_cat_ value we measured for the *P. patens* non-modified mMDH1 in comparison to that of the *A. thaliana* wild-type enzyme (Fig. 4B).

The analysis of the influences of acetylation of K172 in *P. patens,* and K169 and K170 in *A. thaliana,* on the kinetic properties of mMDH1 in the malate oxidation direction indicated a major difference between both plant species. While the kinetic properties of *P. patens* and *A. thaliana* mMDH1 acetylated at the corresponding lysine residues (K172 in *P. patens* and K169 in *A. thaliana*) were not modified, acetylation of *A. thaliana* mMDH1 at K170 reduces the enzymatic catalytic efficiency to 67.5 % of the non-modified enzyme due to a high reduction of the affinity for malate (Fig. 4B).

The lysine residues corresponding to K169 and K170 of *A. thaliana* mMDH1 that influence the enzymatic activity of *A. thaliana* and *P. patens* mMDH1 in an acetylation-dependent manner are conserved in plants and were found to be acetylated in several species. In no case the residues were found acetylated simultaneously. The fact that K169 and K170 were found to be independently acetylated in different organs (leaves and embryos) of rice may indicate the need of fine-tuning the enzyme activity to cope with specific metabolic necessities of the organs.

Our analysis of the modelled protein structures indicates that the critical lysine residues are not near the catalytic site or in the dimer or tetramer interfaces. While the precise molecular mechanism of how acetylation influences the catalytic properties of the enzyme remains to be resolved, it appears likely that acetylation induces a conformational change that reaches across the protein structure.

### Acetylation of a conserved lysine at the carboxyl terminal end of *A. thaliana* mMDH1 favors the malate oxidation activity

The C-terminus of *A. thaliana* mMDH1 contains three lysines (K325, K329, and K334) that were found acetylated in seedlings harvested at the beginning of the light period (König, 2014). K334 is conserved in all plant species investigated in this study, while K325 and K329 are shared by ~80% and ~70% of the investigated plant species, respectively (Fig. 3). The conservation of these lysine residues in plants suggests a possible conservation of their regulatory roles; yet to date, these residues were only found acetylated in mMDH orthologs of a few species beyond *A. thaliana* (Fig. 3, Supporting Table S1). We found that acetylation at the single positions K325 or K329 slightly influence the catalytic behavior of mMDH1. In contrast, acetylation at the highly conserved K334 has a major impact on the malate oxidative activity: compared to the non-modified enzyme, it decreases the catalytic efficiency of OAA reduction to 40% while increasing the catalytic efficiency of malate oxidation by 3.4-fold. A similar increase of the catalytic efficiency of the malate oxidation reaction was reported as a consequence of acetylation of MDH in other organisms. Acetylation of *E. coli* MDH at K140, which is located next to *A. thaliana* K170 and *P. patens* K172 (Fig. 6C), was shown to increase the catalytic efficiency of malate oxidation by 3.4-fold (Venkat *et al.*, 2017). In the same reaction direction, the catalytic efficiency of the *E. coli* enzyme is also enhanced by acetylation of K99 (by 2.9-fold), while that of human mMDH2 is enhanced by acetylation of K307 (by 3.1-fold) (Venkat *et al.*, 2017). However, different to Arabidopsis, the changes in the catalytic efficiency observed in *E. coli* and human MDH were due to increases of the enzymatic activity, while the affinity for malate was unchanged. This indicates that the activation of mMDH in different organisms involves alternative molecular mechanisms.

### Potential consequences of mMDH lysine acetylation for plant metabolism

Our results indicate that acetylation of *A. thaliana* mMDH1 at the single lysine position K170 reduces the efficiency of the reaction in both directions by a similar value (67 % efficiency in the OAA reduction and 72 % efficiency in the malate oxidation direction). Moreover, we found that acetylation at K334 favors the malate oxidation activity of *A. thaliana* mMDH1, as it decreases the efficiency of OAA reduction while increasing the efficiency of malate oxidation. Since acetylation of K170 and K334 allow adjustments in both directions of the reaction catalysed by mMDH1, which are fully reversible and do not require protein synthesis and/or turnover, they likely represent effective mechanisms to modulate flux through central carbon metabolism. Knowing the stoichiometry of acetylation could allow a more precise description of the impact of acetylation of a protein in metabolism (Weinert *et al.*, 2014), even though low PTM site stoichiometry does not indicate a minor or lack of function (Xia *et al.*, 2012, Bovdilova *et al.*, 2019, Hansen *et al.*, 2019). The degree of acetylation of the individual lysines analysed here is likely to vary *in vivo* depending on conditions and importantly on the cellular context. While PTM stoichiometry in a total tissue extract is a poor predictor of *in vivo* significance, the finding that acetylation of the same conserved lysine residues occurs in different plant species, even under different conditions, is an indicator of the functional relevance of the sites.

The results of the kinetic analysis of the individual acetylated sites may inform hypotheses about possible metabolic contexts in which the analysed acetylation sites of plant mMDH may be relevant. Under conditions of high NAD^+^/NADH ratio in the matrix, the activity of mMDH1 in the OAA reduction direction needs to be limited, while that of the malate oxidation boosted to avoid further removal of reductant from the matrix and constraints on mitochondrial ATP production in turn (Fig. 1). Acetylation of *P. patens* mMDH1 at K172 increases the capacity – and potentially the rate – of OAA consumption, which will be necessary under metabolic conditions that demand a high rate of matrix NAD^+^ provision at the expense of ATP production, e.g., during photorespiratory conditions (Fig. 1). In *E. coli*, the activity of MDH in the malate oxidation direction and the acetylation grade of the enzyme increase with increasing glucose concentrations (Venkat *et al.*, 2017), suggesting an activation of the enzyme to provide sufficient capacity for enhanced flux through the TCA-cycle in its circular mode. Interestingly, bacteria grown on citrate, which favors the OAA reduction reaction of MDH, showed lower acetylation of MDH compared to growth on glucose (Wang *et al.*, 2010, Mall *et al.*, 2018).

The lysine residues found to be acetylated in mMDH1 of *A. thaliana* and *P. patens* most likely represent only a small proportion of the mMDH1 lysine sites that can be acetylated *in vivo*. As PTMs can be strictly dependent on exact conditions, other mMDH lysine residues may also be acetylated under different growth conditions, at different times of the day, or in other tissues. Additional, systematic analyses will be required in the future for their identification. Nevertheless, as in the case of other PTMs, not all acetylated lysines necessarily modify the kinetic parameters of an enzyme, as we here observed in some cases (Fig. 4B and C). Alternatively, these modifications may influence interactions with other proteins or factors that are not present in the *in vitro* setting devised here. Indeed, *in vivo* interactions between several mitochondrial proteins that were found acetylated in this study were recently reported in *A. thaliana in vivo* (Zhang *et al.*, 2018), raising the possibility of further regulatory functions, e.g., by modulating metabolic channeling (Zhang *et al.*, 2017, Zhang *et al.*, 2018).

Our present analysis offers intriguing insights into the functional impact of single-site modifications on the enzymatic kinetics. It further provides evidence for the degrees of potential *in vivo* regulation, which are likely key for the regulation of cellular metabolism but are not usually accounted for by standard reductionist *in vitro* analyses. Acetylation of evolutionary conserved lysines provides a mechanism for tuning enzyme function to specific metabolic requirements. *In vivo*, the exact acetylation pattern across different lysine residues, which may dynamically change with the metabolic status of the cells, will govern the dynamic regulation of mMDH performance. Ideally, novel highly sensitive mass spectrometry-based methods will become available in the future, which will enable us to assess the PTM site occupancies on proteins in single cells and organelles of certain plant tissues. Such analyses would allow the direct identification of relevant PTMs that regulate metabolic activity *in vivo*. Also, the interplay of the different modifications involved in the modulation of mMDH as well as the conditions that lead to the specific changes in mMDH acetylation status of evolutionary conserved lysines remain to be investigated in the future.

## Experimental procedures

### *Physcomitrium patens* cultivation conditions

*P. patens* (Hedw.) Bruch & Schimp. strain Gransden (Rensing *et al.*, 2008) gametophores were cultivated on modified Knop medium plates (KH_2_PO_4_ (250 mg L^−1^), KCl (250 mg L^−1^), MgSO_4_ x 7H_2_0 (250 mg L^−1^), Ca(NO_3_)_2_ x 4H_2_O (1000 mg L^−1^), FeSO_4_ x 7H_2_O (12.5 mg L^−1^), CuSO_4_ (0.22 μM), ZnSO_4_ (0.19 μM), H_3_BO_3_ (10 μM), Na_2_MoO_4_ (0.1 μM), MnCl_2_ (2 μM), CoCl_2_ (0.23 μM) and KI (0.17 μM), pH = 5.8, 1 % [w/v] agar; (Rudinger *et al.*, 2011)) at 2° C under long day conditions with 16 h light and 8 h darkness at a light intensity of 65 μmol m^−2^ s^−1^ using neon tubes Osram HO 39W/865 Lumilux Cool Daylight. Single gametophores were transferred onto fresh plates three weeks prior to harvest and were further cultivated at 10 h light and 14 h darkness.

### Trypsin digestion, immuno-enrichment of lysine-acetylated peptides and LC-MS/MS data acquisition

The lysine acetylome analysis was performed as described in detail in Lassowskat et al. (2017). Briefly, proteins were extracted from three independent replicates (including 5-6 gametophores/plate and 9-15 plates per replicate). Protein extracts were alkylated and digested with MS-grade trypsin. Peptides were desalted and enriched for lysine-acetylated proteins by immuno-purification. Total peptide samples and the immuno-enriched fractions were then analyzed via LC-MS/MS using an EASY-nLC 1200 (Thermo Fisher) coupled to a Q Exactive HF mass spectrometer (Thermo Fisher). Peptides were separated on 20 cm frit-less silica emitters (New Objective, 0.75 μm inner diameter), packed in-house with reversed-phase ReproSil-Pur C18 AQ 1.9 μm resin (Dr. Maisch). The column was kept at 50°C in a column oven throughout the run. The following parameters were used for acetylome analysis: Peptides were eluted for 78 min using a segmented linear gradient of 0 to 98 % solvent B (solvent A 0 % ACN, 0.1 % FA; solvent B 80 % ACN, 0.1 % FA) at a flow-rate of 250 nL/min. Mass spectra were acquired in data-dependent acquisition mode with a Top12 method. MS spectra were acquired in the Orbitrap analyzer with a mass range of 300–1759 m/z at a resolution of 120000 FWHM, maximum IT of 55 ms and a target value of 3 × 10^6^ ions. Precursors were selected with an isolation window of 1.2 m/z. HCD fragmentation was performed at a normalized collision energy of 25. MS/MS spectra were acquired with a target value of 5 × 10^4^ ions at a resolution of 15000 FWHM, maximum IT of 150 ms and a fixed first mass of m/z 100. Peptides with a charge of +1, greater than 6, or with unassigned charge state were excluded from fragmentation for MS2, and dynamic exclusion for 30 s prevented repeated selection of precursors.

### MS data analysis

Raw data were processed using the MaxQuant software V1.6.17 (http://www.maxquant.org/) (Cox and Mann, 2008). MS/MS spectra were searched with the Andromeda engine against the cosmoss_V3.3 database (February 15, 2017, https://www.cosmoss.org/physcome_project/wiki/Downloads). Sequences of 248 common contaminant proteins and decoy sequences were automatically added during the search. Trypsin specificity was required and a maximum of four missed cleavages were allowed. Minimal peptide length was set to seven amino acids. Carbamidomethylation of cysteine residues was set as fixed, oxidation of methionine and protein N-terminal acetylation, acetylation of lysines as variable modifications. Light and medium dimethylation of lysines and peptide N-termini were set as labels. Peptide-spectrum-matches and proteins were retained if they were below a false discovery rate of 1 %, modified peptides were additionally filtered for a score ≥ 35 and a delta score of ≥ 6 to remove low quality identifications. Match between runs and requantify options were enabled. For acetylome analyses reverse hits and contaminants were removed. Acetylation sites were filtered for a localization probability of ≥ 0.75.

### MS analyses of the recombinant mMDH proteins

The recombinant proteins (10 μg of each) were alkylated and trypsin-digested in urea buffer as described previously (König *et al.*, 2014b). Up to 500 ng of the trypsinated peptides were analysed by LC-MS/MS as described above with the following differences: Mass spectra were acquired in data-dependent acquisition mode with a Top15 method. MS spectra were acquired in the Orbitrap analyzer with a mass range of 300–1759 m/z at a resolution of 60000 FWHM, maximum IT of 30 ms and a target value of 3 × 10^6^ ions. Precursors were selected with an isolation window of 1.3 m/z. MS/MS spectra were acquired with a target value of 1 × 10^5^ ions at a resolution of 15000 FWHM, maximum IT of 55 ms. For the data analysis, MaxQuant software was used with standard settings and the following additions: MS/MS spectra were searched with the Andromeda engine against the Uniprot *E. coli* k12 database including either the protein sequences of *P. patens* or *A. thaliana* mMDH. Trypsin specificity was required and a maximum of two missed cleavages were allowed. Carbamidomethylation of cysteine residues was set as fixed, oxidation of methionine and acetylation of lysines as variable modifications. Results of all mMDH peptides from modification specific peptides table are presented in Supporting Table S6. It has to be noted that some acetylation sites are present on the recombinant mMDH proteins due to untargeted acetylation from *E. coli*, however these acetylation sites are only present at very low level (mostly below 1%) as estimated from the peptide intensities compared to the unmodified peptide intensities.

### Functional classification and prediction of subcellular localization

Proteins were functional annotated based on the *P. paten*s V3.3 database. Localization was inferred from the experimental dataset by Müller et al. (2014) and the GO_CC annotation provided in Lang et al. (2018).

### Comparison of acetylated lysines of putative mitochondrial orthologs

Sequences of the 120 acetylated *A. thaliana* proteins identified in König et al. (2014a) were obtained from Tair (https://www.arabidopsis.org/tools/bulk/sequences/index.jsp) and sequences of those *P. patens* proteins that were acetylated and classified were obtained from Peatmoss (https://peatmoss.online.uni-marburg.de/ppatens_db/pp_search_input.php) (Fernandez-Pozo *et al.*, 2020). Amino acid sequences of both species were combined and clustered using the multiple alignment program MAFFT Version 7 with default settings (Katoh *et al.*, 2002, Katoh *et al.*, 2019). *P. patens* and *A. thaliana* proteins sequences of orthologous groups were aligned again with Muscle (Edgar, 2004) using the Mega 7 software (Kumar *et al.*, 2016). In total 22 clusters of *P. patens* and *A. thaliana* proteins were identified with homologs sharing sequence similarity higher than 40 %. In each cluster different numbers of paralogous proteins of each species were included. Acetylation sites and amino acid positions of proteins of each orthologous group were compared. Acetylation sites were listed and positions extracted from the original alignment (Table 1, Supporting Table S4, Supporting Data S1).

### Phylogenetic analysis, comparison of sequence conservation and lysine acetylation patterns of mMDHs of different plant species

Published acetylome datasets were inspected for the presence of the mitochondrial malate dehydrogenase. In total 17 datasets for 10 different flowering plants were included in the analysis (Supporting Table S5). For seven flowering plant species lysine acetylation of mMDH proteins was detected. Plant tissues or organs, in which these lysine acetylated mMDHs were detected, are given in Supporting Table S5. Putative mMDH orthologs of these species were identified by Basic Local Alignment Tool (BLAST) searches using the *A. thaliana* mMDH1 protein sequence as query against nonredundant protein database (BLASTP) and against translated nucleotide database (TBLASTN) (Altschul *et al.*, 1990) of different sources. Sequence data of the selected angiosperm species *C. sinensis*, *V. vinifera*, *O. sativa, B. distachyon, F. ananass* and the green algae *C. reinhardii* were deposited at NCBI (www.ncbi.nlm.nih.gov) and sequence data of *P. patens* and *M. polymorpha* are deposited at Phytozome v12.0 (https://phytozome.jgi.doe.gov/pz/portal.html). For putative mMDH orthologs of Azolla and Salvinia, sequences are available in the Fernbase (www.fernbase.org). For *P. sativum* the URGI database was used as sequence source (Kreplak *et al.*, 2019). The *Anthoceros* sequences were retrieved from the recently published *A. agrestis*, Bonn v1 strain genome v1.1 (Li *et al.*, 2020). Best 2-10 hits were downloaded for each species and were aligned in the MEGA alignment explorer with the MUSCLE tool (Tamura *et al.*, 2015, Kumar *et al.*, 2016) followed by manual adjustment.

Amino acid sequence alignment was used to calculate a Neighbor joining phylogeny with 1000 bootstraps (Poisson model, gamma distributed, partial deletion with cut off 90%) including the peroxisomal (AT2G22780), plastidial (AT3G47520) and mitochondrial *A. thaliana* mMDH proteins. With a reduced dataset (Supporting Data S2) including all MDH proteins of the identified mitochondrial cluster, glyoxysomal/peroxisomal and plastid MDHs of *P. patens* and *A. thaliana* and the MDH of *E. coli* and the mMDH2 of human, we constructed a Maximum Likelihood phylogram using the IQ-tree software version 1.6.12 (Trifinopoulos *et al.*, 2016). We applied the LG+G4 model for sequence evolution as the best fitting model identified by ModelFinder (Kalyaanamoorthy *et al.*, 2017). We determined the confidence of each node from 1,000 bootstrap replicates with ultrafast bootstrap approximation “UFBoot” (Hoang *et al.*, 2018). In parallel we calculated a Neighbor joining phylogeny with same settings as described above (Supporting Fig. S1). Sequences of mMDH proteins identified to be acetylated were aligned and conservation was displayed using the GeneDoc software (https://genedoc.software.informer.com/2.7/).

### Cloning of *A. thaliana* mMDH1 into the expression vector

The cDNA encoding the mMDH1 mature protein (without signal peptide) from *A. thaliana* (Hüdig *et al.*, 2015) was introduced in pCDF PylT-1 vector (SmR; *T7* promoter; *T7* terminator) (Neumann *et al.*, 2009). This vector allows the expression of proteins fused to an N-terminal His tag for purification by Ni-NTA affinity chromatography. The mMDH cassette was obtained by PCR using primers mmdh1_GibFw and mmdh1_GibRv and cloned by Gibson assembly. The sequence of the resulting plasmid was verified by sequencing. The pCDFmdh1 vector was used as template for site directed mutagenesis by PCR, to change specific lysine codons to the amber codon, using the primers listed in Supporting Table S6. The following constructs were generated: pCDFmdh1amb170, pCDFmdh1amb325, pCDFmdh1amb329, pCDFmdh1amb334. The incorporation of the desired nucleotide changes was confirmed by sequencing (Macrogen).

### Cloning of *P. patens* mMDH1 into the expression vector

The coding sequence of mMDH1 (without signal peptide) of *P. patens* Gransden (gene Pp1s201_6V6.1) (Lang *et al.*, 2018) was amplified with primers MfeI_PpMDH1_f and KpnI_PpMDH1_r (Supporting Table S6). The mMDH1 version with the amber codon for amino acid position 172 was generated by overlap extension PCR with internal primers (PpMDH1_KStop_f and PpMDH1_KStop_r, Supporting Table S6). The fragments were introduced into the pCDF PylT-1 vector (SmR; *T7* promoter; *T7* terminator) between restriction sites *MfeI and KpnI* (SmR; *T7* promoter; *T7* terminator) (Neumann *et al.*, 2009) to generate N-terminally His-tagged mMDH1. The incorporation of the desired nucleotide changes was confirmed by sequencing (Macrogen).

### Expression and purification of acetylated mMDHs

All pCDF PylT expression constructs carrying the mMDH1 versions with an amber codon instead of the codon for the lysine to be acetylated and those carrying the corresponding wild-type mMDH1 were transformed into *E. coli* Rosetta2 (DE3). Plasmid pBK-AcKRS3, carrying the acetyl-lysyl-tRNA synthetase (AcKRS) (Neumann *et al.*, 2008), was co-transformed with the pCDF PylT plasmids. For heterologous protein production, transformed cells were grown in 400 ml LB medium in the presence of the appropriate antibiotic in each case at 37° C and 16 *g* to an OD600 of 0.4 - 0.6. The growth medium was supplemented with 10 mM N -acetyl-lysine and 20 mM nicotinamide 30 min before induction. To induce the expression of the heterologous protein, 1 mM IPTG was added to the culture and the cells were incubated for 20 h at 30° C and 16 *g*. The cellular culture was harvested at 6,000 *g* for 15 min, and pellets were stored at −20° C until use.

For protein extraction, pellets were thawed on ice, resuspended in 20 mM Tris-HCl (pH 8.0) containing 500 mM NaCl, 5 mM imidazole, 1 mM phenylmethylsulfonyl fluoride and a spatula-tip of lysozyme, sonicated and centrifuged at 14,000 *g* for 20 min at 4° C. The supernatant was used for protein purification by gravity-flow immobilized metal ion chromatography on nickel-nitrilotriacetic acid agarose (Ni-NTA Agarose, ThermoFisher). The column was pre-equilibrated with 20 mM Tris-HCl buffer containing 500 mM NaCl and 5 mM imidazole. After loading of the supernatant, the columns were washed in four steps with 500 mM NaCl in 20 mM Tris-HCl (pH 8.0) containing increasing concentrations of imidazole (5, 40 and 60 mM). Protein elution was performed in four steps of 500 μl of 20 mM Tris-HCl, 500 mM NaCl and 200 mM imidazole. Finally, Vivaspin® 10K columns (Satorius, Germany) were used for protein concentration and buffer exchanging (HEPES 50 mM, pH 7.4). The isolated recombinant proteins were analysed by SDS-PAGE and LC-MS/MS (see above) to verify integrity and purity. Protein concentration was determined by Amido black assay (Dieckmann-Schuppert and Schnittler, 1997).

### Kinetic characterization of acetylated mMDH1

MDH activity assays were carried on in a Synergy HT Biotek Plate Reader system at 25° C. OAA reduction was determined following the oxidation of NADH at 340 nm (ε= 6.2 cm^−1^ mM^−1^). The assay medium contained 50 mM Hepes-KOH pH 7.5, 10 mM MgCl_2_, 5 mM NADH and variable concentrations of OAA in the range 0.01 to 4 mM. Malate oxidation was determined following the reduction of NAD at 340 nm in an assay medium containing 50 mM Hepes-KOH pH 7.5, 10 mM MgCl_2,_ 4 mM NAD and a variable concentration of malate in the range 0.01 to 10 mM.

The reactions were started by the addition of the substrate. The kinetic constants were calculated with a minimum of three different enzyme batches each in at least triplicate determinations and adjusted to Michaelis-Menten equation by non-linear regression with Prism 6 (GraphPad Software).

### Gel electrophoresis

SDS-PAGE was performed using 14% (w/v) polyacrylamide gels according to (Laemmli, 1970) and the molecular weight markers Spectra Multicolor Broad Range Protein Ladder from ThermoFisher Scientific. Proteins were visualized by staining with Coomassie Brilliant Blue. Purified enzymes were analyzed on a non-denaturing 7 % (w/v) polyacrylamide gel using the markers ß-amlyase (200 kDa), alcohol dehydrogenase (150 kDa) and bovine serum albumin (66 kDa) from Sigma Aldrich. In-gel mMDH activity assays were performed by incubating the gels in the dark at room temperature in a reaction buffer containing 50 mM K_2_PO_4_ (pH 7.4), 5 mM malate, 0.2 mM NAD, 0.05 % (w/v) NBT and 10 μM PMS as described in Hüdig et al. (2015).

### Statistical analyses

Statistical analyses of the kinetic data were performed on data from at least three biological replicates, each measured at least in triplicate. To verify the statistical differences, P values were determined using unpaired t-test. Statistical analyses of the distribution of acetylation in organellar proteins and non-organellar proteins were performed with chi-square test using Excel package, XLSTAT (Addinsoft).

### Homology modelling of mMDH 3-D structures

The 3-D models of *E. coli* MDH (PDB code 1EMD) and *H. sapiens* MDH2 (PDB code 2DFD) structures were obtained from the RSCB Protein Data Bank (https://www.rcsb.org/). Both *A. thaliana* and *P. patens* mMDH1 models were obtained by using the phyre2 protein structure prediction server (http://www.sbg.bio.ic.ac.uk/~phyre2/html/page.cgi?id=index) with the corresponding amino acid sequence as input. The ligand molecules NAD^+^ and citrate were manually fitted in by aligning the structure of the *H. sapiens* MDH2 (pdb code 2DFD) to the modelled structures of *A. thaliana* and *P. patens* mMDH1. Images of the models were prepared using PyMOL v.2.4.0 by Schrödinger (https://pymol.org/).

## Data statement

The data underlying this article are available in the article and in its online Supporting material.

## Author contributions

M.B., M.M.B., A.B., M.H., and L.R. performed biochemical studies on recombinant proteins. M.E. and J.G. generated the lysine acetylome of *P. patens*. M.S., I.F, M.S-R, and V.G.M. designed the experiments and analyzed the data together with other researchers; I.F, M.S-R, and V.G.M. conceived the project. I.F, M.S-R, and V.G.M. wrote the article with contributions of M.S., M.B. and M.E.; V.G.M. agrees to serve as the author responsible for contact and ensures communication.

## Acknowledgements

This work was enabled through the collaborative DFG research grant PAK918 (IF1655/3-1, MA2379/14-1, SCHA1953/3-1, and SCHW1719/5-1) as part of the ‘Plant Mitochondria in New Light’ initiative and INST 211/744-1 FUGG for LC-MS/MS analyses. Work of MBB in the group of VGM was funded by the StayConnected-Grant of the Heinrich Heine University Düsseldorf. The pBK PylS and pCDF PylT vectors were provided by Dr. J. Chin (University of Cambridge) under a non-commercial material transfer agreement to IF. We thank Astrid Höppner (HHU, Düsseldorf) for her help with the structural analysis and Paulina Pieloch and Dr. Jürgen Eirich for technical assistance within the MS Proteomics Unit Biology of Plants of the University of Muenster. The co-authors have no conflict of interest to declare.

## Supporting material

Supporting Data S1. Protein alignment of identified orthologous groups.

Supporting Data S2. Sequence alignment of the MDH dataset for phylogenetic construction.

Supporting Fig. S1. Mitochondrial malate dehydrogenases along land plant phylogeny.

Supporting Table S1. Proteins identified in total extracts from Physcomitrella gametophores, which were immuno-enriched for lysine-acetylated peptides.

Supporting Table S2. Lysine-acetylated proteins identified in total extracts from Physcomitrella gametophores after immuno-enrichment of lysine-acetylated peptides.

Supporting Table S3. Acetylated lysine sites

Supporting Table S4. Conservation of lysine acetylation between mitochondrial proteins of orthologous groups identified in A. thaliana and P. patens.

Supporting Table S5. Lysine acetylome data of mMDH proteins in plants, human and *E. coli*

Supporting Table S6. Unique peptides of mMDH recombinant proteins purified from *E. coli*.

Supporting Table S7. Primers used in this work.

## Notes

### Competing Interest Statement

The authors have declared no competing interest.

