## Supplementary material for "Acetylation of conserved lysines fine-tune mitochondrial malate dehydrogenase activity in land plants": Suppl Fig S1

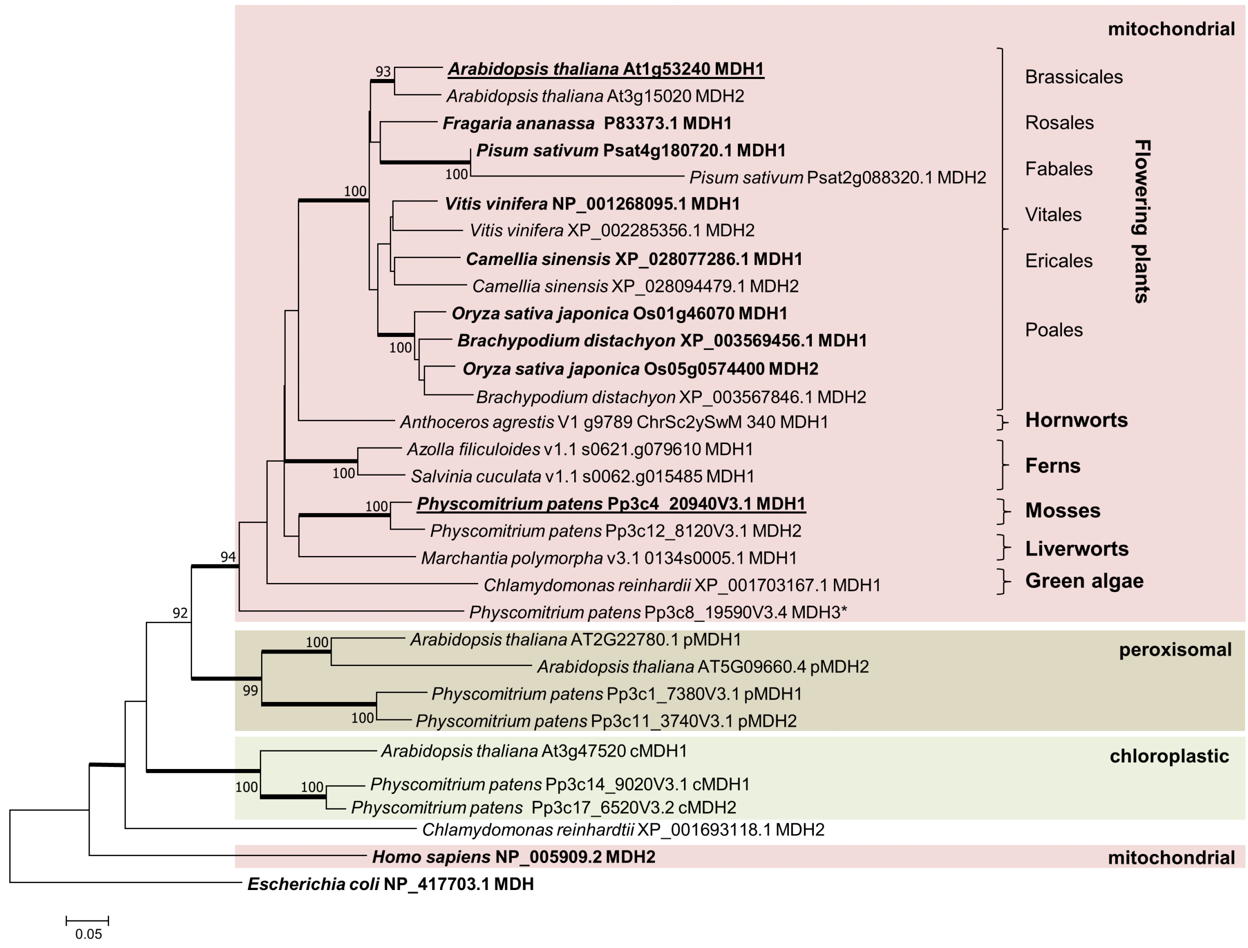


**Supporting Figure S1. Mitochondrial malate dehydrogenases along land plant phylogeny.** Homologs of *A. thaliana* mMDH1 protein were extracted from available sequence data of model species of the main land plant clades. mMDHs of plant species with information on their lysine acetylation status as well as chloroplastic (cMDH) and peroxisomal (pMDH) MDHs of *A. thaliana* and *P. patens* are included. The human mMDH2 and *E. coli* MDH were used as outgroup to root the Neighbor joining tree with 1000 Bootstrap iterations. Bootstrap values are shown where exceeding 90. Thickened internode lines indicate bootstrap supports of 99-100 % in the Maximum likelyhood analysis calculated in parallel. For most species two paralogs of mMDHs were identified and labeled as MDH1 (dominant isoform) and MDH2. Isoforms for which acetylation sites were identified to date are labeled in bold, MDH proteins which are objects of this study are underlined. Red box: mitochondrial MDHs (mMDH), green box: chloroplastic MDHs (cMDH), brown box: peroxisomal MDHs (pMDH). For all proteins, species name is followed by accession number of the sequence or gene name (if accessible) and abbreviation. * marks a third MDH paralog (MDH3) encoded in the *P. patens* genome, which was not identified in the available proteomic datasets ((Müller et al., 2014) and this study). Analysis of sequence conservation and expression profiles suggest that MDH3 is likely an unfunctional protein (PEATmoos.org, (Fernandez-Pozo et al., 2020).
