## Supplementary material for "Acetylation of conserved lysines fine-tune mitochondrial malate dehydrogenase activity in land plants": Suppl Table S5

### Supplementary Table 5. Lysine acetylome data of mMDH proteins in plants, human and *E. coli*

| Species | Sequence information | name | Tissue/organ | acK | acK (AtMDH1) | Reference |
|---|---|---|---|---|---|---|
| ***Arabidopsis thaliana*** | At1g53240, Q9ZP06 | MDH1 | isolated mitochondria of seedlings | 170,325,329,334 | **170,325,329,334** | (König et al. 2014) |
| | | | hda14/ WT leaves | identified, not acetylated | | {Hartl, 2017 |
| | | | seedlings, roots, rosettes, flowers, siliques | not identified | | (Uhrig et al. 2019) |
| | | | | 170,325,329,334 | **170,325,329,334** | all studies combined |
| | At3g15020, Q9LKA3 | MDH2 | isolated mitochondria of seedlings | 170* | **170*** | (König et al. 2014) |
| ***Fragaria ananassa*** | P83373.1, P83373 | MDH1 | leaves | 180, 167 | **170, 183** | (Fang et al. 2015) |
| ***Pisum sativum*** | Psat4g180720.1 URGI | MDH1 | isolated mitochondria of seedlings | 138, 164 | 135, **161** | (Smith-Hammond et al. 2014) |
| | Psat2g088320.1 URGI | MDH2 | | | | |
| ***Glycine max*** | | | seedling leaves | not identified | | (Smith-Hammond et al. 2014) |
| ***Vitis vinifera*** | NP_001268095.1, F6HM78 | MDH1 | fruit exocarp | 39, 58, 135, 169, 178, 274, 291, 298 | 31, 50, 127, **161**, **170**, **301**, 318, **325** | (Melo-Braga et al. 2012) |
| | | | fruit mesocarp | 39, 58 | 31, 50 | |
| | XP_002285356.1 | MDH2 | | | | |
| ***Solanum tuberosum*** | | | isolated mitochondria of tubers | not identified | | (Salvato et al. 2014) |
| ***Camellia sinensis*** | XP_028077286.1 | MDH1 | hda14/ WT leaves | 173, 187, 193 | **169, 183**, 189 | (Jiang et al. 2018) |
| | XP_028094479.1 | MDH2 | | | | |
| ***Oryza sativa ssp. japonicum*** | Os01g46070, XP_015621604.1, Q94JA2 | MDH1 | seed embryos | 333 | **334** | (He et al. 2016) |
| | Os05g0574400, XP_015639465.1 Q6F361 | MDH2 | seed embryos | 167, 180, 327, 335 | **170, 183, 329**, 337 | (He et al. 2016) |
| | | | seedling leaves | 166,180, 327 | **169, 183, 329** | {Zhou, 2018) |
| | | | rice plant | not identified | | (Nallamilli et al. 2014) |
| | | | pistil/developing seeds | not identified | | (Wang et al. 2017) |
| | | | complete rice plant | 335 | 337 | (Xiong et al. 2016) |
| | | | | 166, 167, 180, 327, 335 | **169,170, 183, 329**, 337 | all studies combined |
| ***Brachypodium distachyon (Bd21)*** | XP_003569456.1 | MDH1 | seedling leaves | 167 | **169** | (Zhang et al. 2015; Zhen et al. 2016) |
| | XP_003567846.1 | MDH2 | | 167* | **169*** | |
| ***Triticum sativum*** | | | seedling leaves | not identified | | (Zhang et al. 2016) |
| ***Physcomitrium patens*** | Pp3c4_20940V3.1 Phytozome | MDH1 | gametophore | 172 | **169** | this study |
| | Pp3c12_8120V3.1 Phytozome | MDH2 | gametophore | 170* | **169*** | this study |
| | Pp3c8_19590V3.4 Phytozome | MDH3 | gametophore | not identified | | |
| ***Homo sapiens*** | NP_005909.2, P40926 | MDH2 | | 185, 301, 307, 314 | **190**, 306, 312, | (Zhao et al. |

| | | | | | | |
|---|---|---|---|---|---|---|
| | | | | | 319 | 2010) |
| ***Escherichia coli*** | NP_417703.2, P61889 | MDH | *Escherichia coli*, different strains | 54,56, 99, 107, 111, 133, 140, 162, 204, 272, 279, 300, 301 | 81, 83, 127, 135, 139, 161, 168, **190**, 233, 301, 308, **329**, 330 | (Schilling et al. 2015) |
| | | | *Escherichia coli* O139:H28 (strain E24377A / ETEC) | 54,56,111,204,300 | 81, 83, 139, 233, **329** | (Kuhn et al. 2014) |
| | | | *Escherichia coli* O127:H6 (strain E2348/69 / EPEC) | 99, 107,133, 140, 162, 272, 279, 301 | 127, 135, 161, 168, **190**, **301**, 308, 330 | (Kuhn et al. 2014) |
| | | | *Escherichia coli* | 99, 140, 162, 272, 300, 301 | 127, 161, **190**, **301**, **329**, 330 | (Bienvenut et al. 2020) |
| | | | | 54,56, 99, 107, 111, 133, 140, 162, 204, 272, 279, 300, 301 | 81, 83, 127, 135, 139, 161, 168, **190**, 233, **301**, 308, **329**, 330 | all studies combined |

**Note:** Listed are species with lysine acetylome data published (reference) and the tissue, the proteins were isolated from. Sequence information (NCBI accession number and UniProt ID, if not stated differently) and names, used in this study, are given for mMDH proteins of species, for which at least one mMDH ortholog was identified to be acetylated (name is underlined). Acetylation sites are numbered based on their positions in the individual proteins (acK) and based on the corresponding position in the *A. thaliana* MDH1 protein (acK (AtMDH1)). Acetylation site positions which are shared between two or more species are displayed in bold and underlined. *: Same peptide as in MDH1.
