## Supplementary material for "Acetylation of conserved lysines fine-tune mitochondrial malate dehydrogenase activity in land plants": Suppl Table S7

**Supplementary Table 6. Primers used in this work.**

| **Name** | **Sequence** |
| --- | --- |
| mmdh1_GibFw | 5´TACCATGGGCAGCAGCCATCACCATCATCACCAACAGCCAGGATCCGGAAAGTCGCCATCCTTGG-3´ |
| mmdh1_GibRv | 5´GCAAGCTTGTCGACCTGCAGGCGCGCCGAGCTCGAATTCGTCACTGGTTGGCAAACTTG-3´ |
| mmdh1amb170_fw | 5´- TAGTTGTTTGGTGTTACCACTCTTG-3´ |
| pET16b_mmdh1_170_rv | 5´- CTTTTCATCGTACATACCAGCCTTC-3´ |
| mdh1amb325_fw | 5´- TAGCCAGAACTCAAGTCCTCC-3´ |
| pET16b_mmdh1_325_rv | 5´- CAATGCTTCCAAGCCTTCCTTC-3´ |
| mdh1amb329_fw | 5´- TAGTCCTCCATAGAAAAGGGAGTC-3´ |
| pET16b_mmdh1_329_rv | 5´-GAGTTCTGGCTTCAATGCTTCC-3´ |
| mmdh1amb325+newF | 5´- TTGTAGCCAGAACTCTAGTCCTC-3´ |
| mmdh1amb325+newR | 5´- TGGCTACAATGCTTCCAAGCCT-3´ |
| mmdh1amb329+newF | 5´- CAGAACTCTAGTCCTCCATAGAATAGG-3´ |
| mmdh1amb329+newR | 5´- GACTAGAGTTCTGGCTTCAATGC-3´ |
| mmdh1amb334newF | 5´- TAGAATAGGGAGTCAAGTTTGCCAAC-3´ |
| mmdh1amb334newR  mdh1amb169_fw  mdh1amb169_rv | 5´- GACTCCCTATTCTATGGAGGACTTG-3´  5´- GTACGATGAATAGAAATTGTTTGG-3´  5´- CCAAACAATTTCTATTCATCGTAC-3´ |
| MfeI_PpMDH1_f | 5´- GGGCAATTGATGTCGTCGCAGACTCCCAAG-3´ |
| KpnI_PpMDH1_r | 5´- CTCGGTACCACTTCAAGCCTTCACATTCACG-3´ |
| PpMDH1_KStop_f | 5´- GGTACTTATGATCCCTAGAGGCTTTTTGGTGTC-3´ |
| PpMDH1_KStop_r | 5´- GACACCAAAAAGCCTCTAGGGATCATAAGTACC-3´ |
| pCDFpylTfor | 5´- TATTGTACACGGCCGCATAATCG-3´ |
| pCDFpylTrev | 5´- CAGCAGCCTAGGTTAATTAAGC-3´ |
| BamHIXaMfeIfor | 5´- GATCCGCATATCGAAGGTCGTC-3´ |
| BamHIXaMfeIrev | 5´- AATTGACGACCTTCGATATGCG-3´ |
